# Functional architecture of neural circuits for leg proprioception in *Drosophila*

**DOI:** 10.1101/2021.05.04.442472

**Authors:** Chenghao Chen, Sweta Agrawal, Brandon Mark, Akira Mamiya, Anne Sustar, Jasper S Phelps, Wei-Chung Allen Lee, Barry J Dickson, Gwyneth M Card, John C Tuthill

**Affiliations:** Department of Physiology and Biophysics, University of Washington, Seattle, WA, USA; Janelia Research Campus, Howard Hughes Medical Institute, Ashburn, VA, USA; Department of Neurobiology, Harvard Medical School, Boston, MA, USA

## Abstract

To effectively control their bodies, animals rely on feedback from proprioceptive mechanosensory neurons. In the *Drosophila* leg, different proprioceptor subtypes monitor joint position, movement direction, and vibration. Here, we investigate how these diverse sensory signals are integrated by central proprioceptive circuits. We find that signals for leg joint position and directional movement converge in second-order neurons, revealing pathways for local feedback control of leg posture. Distinct populations of second-order neurons integrate tibia vibration signals across pairs of legs, suggesting a role in detecting external substrate vibration. In each pathway, the flow of sensory information is dynamically gated and sculpted by inhibition. Overall, our results reveal parallel pathways for processing of internal and external mechanosensory signals, which we propose mediate feedback control of leg movement and vibration sensing, respectively. The existence of a functional connectivity map also provides a resource for interpreting connectomic reconstruction of neural circuits for leg proprioception.

## Introduction

Proprioception, the sense of limb position and movement, plays an indispensable role in motor control by providing continuous sensory feedback to motor circuits in the central nervous system. Proprioception is important for inter-leg coordination during locomotion (Bidaye et al., 2018; Burrows, 1996), stabilization of body posture (Bässler and Büschges, 1998; Zill et al., 2004) and motor learning (Bässler et al., 2007; Takeoka et al., 2014). Loss of limb proprioception impairs locomotion and motor control (Akay et al., 2014; Mendes et al., 2013). Thus, mapping neural circuits that process proprioceptive information is a prerequisite to understanding the role of proprioception in motor flexibility and recovery from injury.

Proprioception relies on mechanosensory neurons embedded in joints and muscles throughout the body, which are referred to as proprioceptors. Different types of proprioceptors detect distinct features of body kinematics. In vertebrates, Golgi tendon organs detect mechanical load on the body, while muscle spindles encode muscle fiber length and contraction velocity (Tuthill and Azim, 2018; Windhorst, 2007). Proprioceptors in invertebrates detect similar features. The three predominant classes of proprioceptors in insects are campaniform sensilla, hair plates, and chordotonal neurons (Tuthill and Wilson, 2016b). Dome-shaped campaniform sensilla encode mechanical load by detecting strain in the cuticle (Zill et al., 2004), hair plates act as joint limit detectors (French and Wong, 1976), and chordotonal neurons detect multiple features of joint kinematics (Burns, 1974; Matheson and Field, 1995). Although they differ in structure, the common functional properties of vertebrate and invertebrate proprioceptors suggest that they have convergently evolved to encode similar mechanical features (Tuthill and Azim, 2018).

Compared to other primary senses, the organization of central circuits for leg proprioception remains poorly understood. Pioneering work in larger insect species, such as the locust (Burrows, 1996) and stick insect (Büschges, 1989) characterized the anatomy and physiology of central proprioceptive neurons. However, most of this prior work relied on sharp-electrode recordings from single neurons, which made it challenging to understand how they operate collectively as a circuit to control behavior. Understanding circuit-level architecture and function is aided by the existence of genetic tools to label, manipulate, and record from identified classes of neurons. Such genetic tools have recently become available for proprioceptive circuits in the *Drosophila* ventral nerve cord (VNC), the invertebrate analog of the spinal cord (Court et al., 2020). An additional advantage of *Drosophila* is the existence of an electron microscopy (EM) volume of the adult VNC (Phelps et al., 2021), which enables synapse-level reconstruction of VNC circuits. Together, the combination of genetic tools and connectomics data provides an opportunity to link connectivity and function of central circuits for leg proprioception.

The largest proprioceptive organ in the *Drosophila* leg is the femoral chordotonal organ (FeCO), which is composed of ∼152 mechanosensory neurons (Kuan et al., 2020) located in the proximal femur and attached to the tibia by a series of tendons (Figure 1A). Calcium imaging has revealed that *Drosophila* FeCO neurons can be divided into three basic subtypes: claw neurons encode tibia position, hook neurons encode movement direction, and club neurons encode bidirectional movement and vibration (Mamiya et al., 2018). The axons of each subtype project to distinct regions of the VNC. This organization suggests that signals from different FeCO subtypes may be processed by separate downstream neurons (Figure 1A, right). However, apart from three specific VNC cell classes (Agrawal et al., 2020), little is known about how information from different FeCO subtypes is integrated by downstream circuits in the *Drosophila* VNC that underlie sensation and guide movement of the leg.

**Figure 1.**
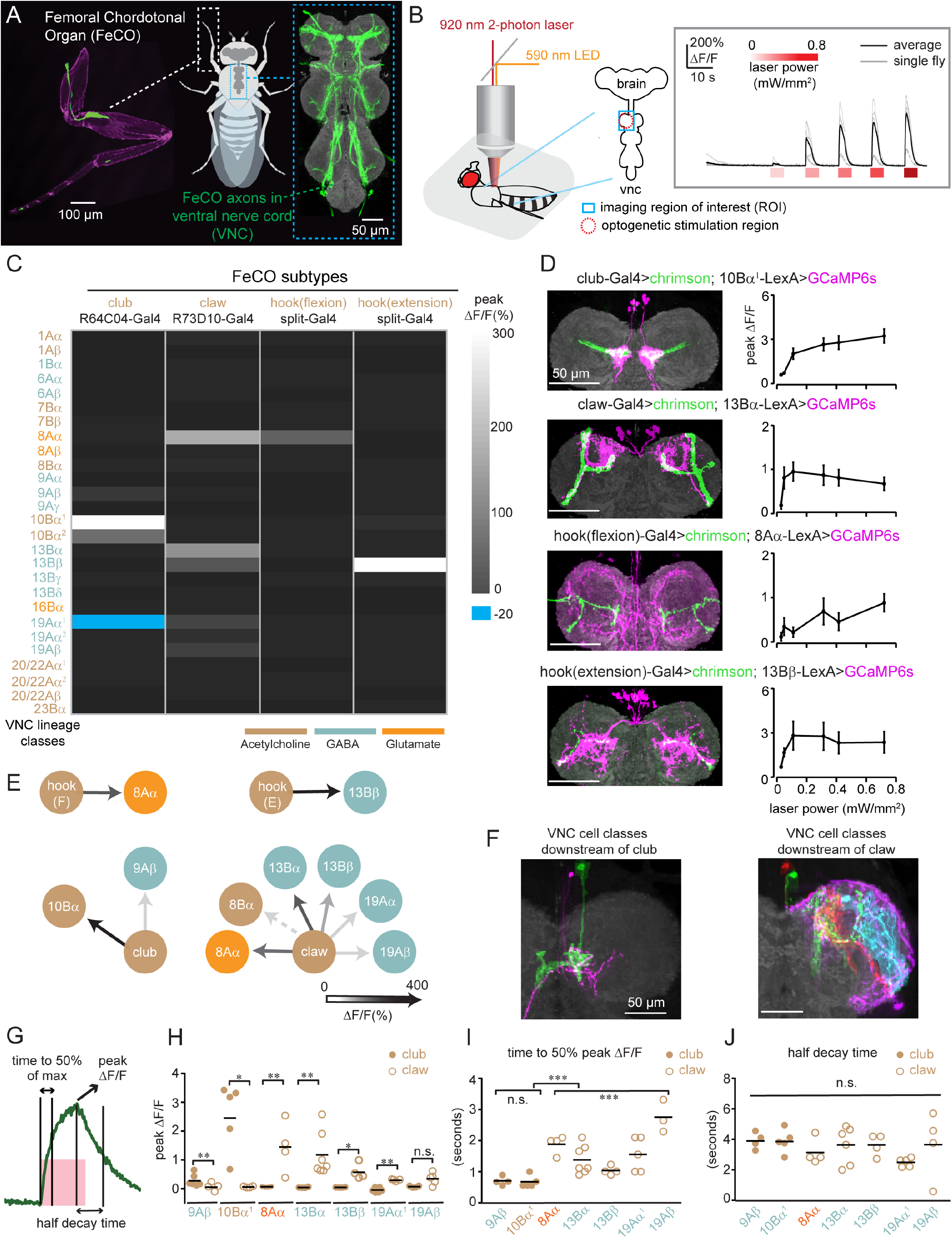
Building a functional connectivity map between FeCO sensory neurons and central neurons in the fly VNC. (**A**) Left: a confocal image of the foreleg (T1) of *Drosophila melanogaster*. The FeCO cell bodies (left) and (axons) are labelled by GFP (green) driven by *iav*-Gal4. Cuticle auto-fluorescence is magenta (left) and the VNC neuropil stained by nc82 is shown in grey (right). (**B**) Experimental set-up for two-photon calcium imaging from VNC neurons while optogenetically stimulating FeCO sensory neurons. Left: Schematic of experimental setup. The blue window indicates the imaging region (ROI) and red dashed circle indicates the region of optogenetic stimulation. Right: example traces of GCaMP6s fluorescence in 10Bα^1^ neurons in response to optogenetic activation of club neurons (n=6 flies). The red bars below the traces indicate the 5-second stimulation window and intensity. **(C)** A heatmap summarizing the average peak calcium signal (ΔF/F) in VNC neurons following optogenetic activation of each FeCO subtype (n>=4 flies). The colors for each lineage and FeCO subtype indicate the putative neurotransmitter that they release. Superscript numbers indicate independent LexA lines that label the same lineage; detailed genotypes are listed in Figure S2. (**D**) Anatomy (left) and peak calcium responses (right, mean ± SEM) of each sensory/central neurons pair (n= 6, 7, 4, 5 flies). **(E**) A summary of the predominant targets downstream of each FeCO subtype. Functional connectivity strength is indicated by the shading of the arrow. Note that the presence of some connections between claw and 8Bα neurons varied across flies (Figure S3), while others were consistent. **(F)** Single neuron anatomy from each class downstream of club (left) and claw (right) sensory neurons were aligned to a common VNC template. (**G**) Quantification of calcium response kinetics. The pink window indicates 5 seconds stimulus duration, while green curve is an example calcium trace. (**H**) Peak calcium response (ΔF/F) (* p<0.05, ** p<0.01, n.s. no significant difference, Mann-Whitney test), (**I**) time to 50% of the maximum signal (* p<0.05, ** p<0.01, n.s. no significant difference, Mann-Whitney test and Kruskal-Wallis test), and (**J**) time to 50% decay from the max for neurons downstream of club (solid brown dots) and claw (open brown circles) sensory neurons (n.s. no significant difference, Kruskal-Wallis test). Each point represents data from an individual fly, while bars indicate the average across flies.

In this study, we elucidate the logic of sensory integration within leg proprioceptive circuits of the *Drosophila* VNC. We first combined two-photon calcium imaging of second-order VNC neurons with optogenetic stimulation of specific FeCO subtypes. This strategy, named “functional connectivity”, has previously been used to map the structure of visual (Morimoto et al., 2020) and navigation (Franconville et al., 2018) circuits in *Drosophila*. Our functional connectivity analysis identified separate circuits for processing tibia vibration and position/movement. We further analyzed spatial and multimodal integration in three specific classes of central neurons. Using spatially targeted optogenetic stimulation to map receptive-field structure, we found that each class either integrates sensory information from multiple FeCO subtypes or from the same FeCO subtype across multiple legs. Finally, we find that inhibition sculpts the adaptation dynamics of second-order neurons encoding leg movement and vibration. Our results demonstrate that diverse proprioceptive signals from different sensory neuron subtypes and locations on the body are directly integrated by second-order neurons and reveal separate central pathways for processing of external substrate vibration and internal self-generated leg joint kinematics.

## Results

We began by creating genetic driver lines to specifically manipulate the activity of each FeCO subtype with optogenetics. Using an anatomical screen of existing driver lines (Jenett et al., 2012; Tirian and Dickson, 2017), we created intersectional Split-Gal4 lines that specifically label club, claw, and hook neurons (Figure S1A), which we previously found encode tibia movement/vibration, position, and direction (Mamiya et al., 2018). To measure the proprioceptive tuning of the neurons labeled by each Split-Gal4 line, we used 2-photon calcium imaging while swinging the tibia between flexion and extension (Figure S1C-D). In addition to confirming the proprioceptive encoding of each subtype, these experiments identified a new FeCO subtype which responds to tibia extension in a directionally-tuned manner. The projections of these neurons are slightly different from the flexion-tuned hook neurons (Figure S1D); therefore, we refer to this new FeCO subtype as “hook-extension”.

With improved tools to specifically target FeCO subtypes, we next sought to identify their downstream partners in the VNC. The VNC is composed of about 20,000 neurons that develop from 34 hemilineages (Harris et al., 2015). Neurons within a hemilineage are anatomically (Shepherd et al., 2019) and transcriptionally (Allen et al., 2020; Lacin and Truman, 2016) similar; they also use the same primary neurotransmitter (Lacin et al., 2019). By visually screening a collection of LexA driver lines, we identified 27 LexA lines that sparsely labeled VNC neurons from each hemilineage that anatomically overlapped with FeCO axons (Figure S2). We denote driver lines that label different neuron classes within the same hemilineage using Greek letters, e.g., 9Aα, 9Aβ, etc.

In *Drosophila*, most neurons release one of three canonical neurotransmitters: acetylcholine, GABA, and glutamate (Allen et al., 2020; Bates et al., 2019). Fast excitation is primarily mediated by acetylcholine, while inhibition is mediated by GABA and glutamate. We were able to infer the neurotransmitter released by each VNC neuron class (Figure 1C, E), based on their hemilineage identity (Lacin et al., 2019) .

We imaged calcium signals from each LexA line in the VNC with GCaMP6s (Chen et al., 2013) while optogenetically stimulating the axons of FeCO neurons expressing the ChR2 variant Chrimson (Klapoetke et al., 2014, Figure 1B). To account for differences in response threshold and synaptic strength, we tested a range of stimulation intensities (Figure 1B). Calcium signals evoked by optogenetic stimulation typically plateaued at a stimulus intensity of 0.3 mW/mm^2^ (Figure 1B-D), so we used this stimulus intensity for subsequent group analyses.

Overall, we identified 8 classes of VNC neurons from 6 lineages that responded to optogenetic stimulation of one or more subtypes of FeCO neurons: 8Aα, 8Bα, 9Aβ, 10Bα, 13Bα 13Bβ, 19Aα, 19Aβ (Figure 1C). Of these, 2 neuron classes (9Aβ, 10Bα) responded to stimulation of club neurons, while the remaining 6 classes responded consistently to the stimulation of claw neurons (Figure 1C). In addition to responding to claw neurons, 8Aα and 13Bβ neurons responded to stimulation of hook-flexion and hook-extension neurons, respectively. The amplitude of calcium signals driven by stimulation of a particular FeCO subtype (e.g., club or claw) varied across different downstream neurons (Fig 1H). Neuron classes that responded to stimulation of club and claw axons were non-overlapping: neurons downstream of the club occupy a medial region of the VNC (mVAC), while neurons downstream of the claw arborize more laterally or in the intermediate neuropil (IntNp, Figure 1F). Comparing calcium dynamics of VNC neurons during proprioceptor excitation revealed that VNC neurons downstream of the club had a faster time to peak than those downstream of the claw (Figure 1I). In contrast, the decay of the calcium response of these two groups (club and claw targets) was similar (Figure 1J). These differences may reflect distinct temporal dynamics in pathways that process leg position vs. vibration-related signals.

In summary, our results reveal the first steps of proprioceptive processing downstream of the FeCO. Signals from club (vibration) and claw (position) are routed to different downstream targets, while claw (position) and hook (direction/movement) signals are combined. We also observed interesting differences in the response dynamics of neurons downstream of the club and claw, consistent with their roles in encoding high and low frequencies, respectively.

### 9Aβ, a VNC cell class downstream of club axons, integrates bidirectional movements and vibration signaling from both prothoracic legs

The two cell classes we identified as downstream targets of the club both have contralateral projections that cross the VNC midline. We therefore hypothesized that they integrate club signals across multiple legs.

We first examined connectivity between club axons and 9Aβ neurons, a class of GABAergic interneurons local to each VNC segment (Figure 2A, left). Single 9Aβ neurons densely innervate the ipsilateral neuromere, but also extend a process contralaterally, across the midline (Figure 2A, right). Identification and reconstruction of 9Aβ and club neurons in an electron microscopy (EM) volume of the *Drosophila* VNC (Phelps et al., 2021) revealed the existence of direct synaptic connections between club axons and ipsilateral 9Aβ neurons (Figure 2B). Signal transmission between FeCO and central neurons may be mediated by chemical synapses, electrical gap junctions, or a mixture of the two. Because all FeCO neurons release acetylcholine (Mamiya et al., 2018), we used an antagonist of nicotinic acetylcholine receptors (MLA, 1 µM) to test for the presence of electrical signaling mediated by gap junctions. MLA blocked club-driven calcium signals in 9Aβ (Figure 2C), suggesting that the connection between club and 9Aβ neurons is mediated by acetylcholine receptors.

**Figure 2.**
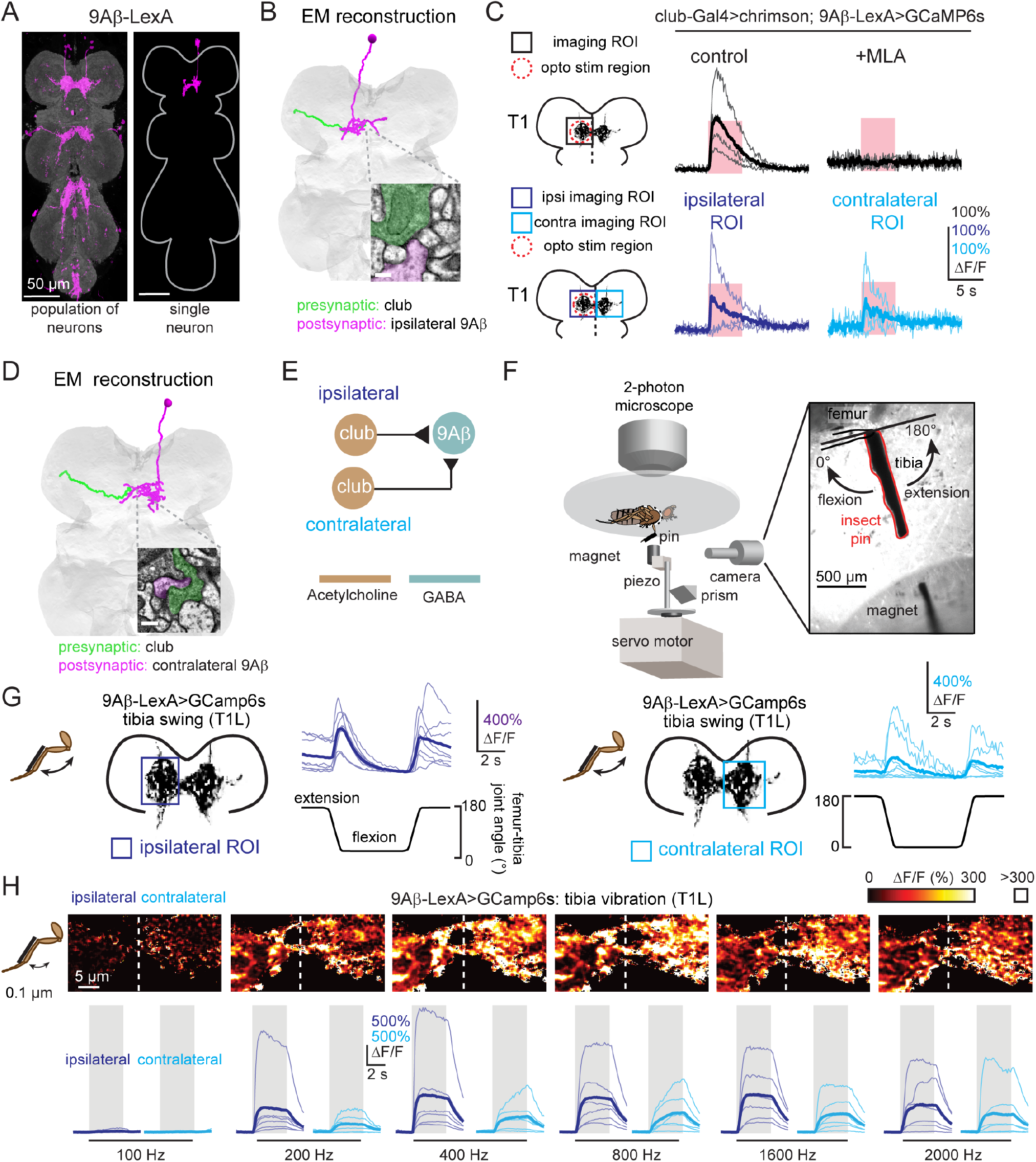
9Aβ neurons receive bidirectional movement and vibration signals from club neurons across both front legs. (**A**) Anatomy of 9Aβ neurons. Magenta is GFP driven by 9Aβ (*R18H03-LexA*), neuropil was stained with nc82 (grey). A single 9Aβ neuron (magenta) is labeled by multi-color FLPout. Both images were aligned to a common VNC template. (**B**) Anatomical reconstruction from EM showing an example of a 9Aβ neuron (magenta) that receives direct synaptic input from an ipsilateral club axon (green). The inset shows an example of a synapse between the two cells. Scale bar = 200nm. (**C**) Calcium response of 9Aβ neurons to optogenetic stimulation of club neurons. Top: calcium responses of 9Aβ neurons in the left prothoracic VNC (T1L) to stimulation of the axons from club neurons in the left foreleg (T1L, n=4). Methyllycaconitine (MLA, 1µM, n=4) effectively blocks excitation from club neurons. Bottom: calcium responses of 9Aβ in left and right neuromeres of the prothoracic VNC (T1L, n=3 and T1R n=3, respectively) to optogenetic stimulation of club axons in T1L (indicated by the red dashed circle). The pink regions indicate stimulus duration (5 seconds, laser power= 0.28 mW/mm^2^). (**D**) Same as in B but showing direct connection between a contralateral 9Aβ neuron (magenta) and a club axon (green) traced from the EM volume. Scale bar = 200nm. (**E**) Proposed wiring diagram for how club axons connect to 9Aβ neurons. (**F**) Experimental setup for calcium imaging during passive leg movements (adapted from Mamiya et al., 2018). Two-photon calcium imaging was used to record calcium signals from the central VNC neurons while controlling and tracking the femur-tibia joint. A pin was glued to the tibia of the front leg and manipulated using a magnet mounted on a motor. The joint was tracked with high-speed video. (**G**) 9Aβ neurons respond to ipsilateral (n=6) and contralateral (n=6) tibia movement. Thin lines are recordings from individual flies; the thicker line indicates the average across flies. (**H**) 9Aβ neurons respond to 0.1 µm vibration of both the ipsi-(n=6) and contralateral (n=6) tibia. Top: The majority of pixels had a ΔF/F value between 0 and 300%; outlier pixels with a value above 300% ΔF/F were set to white for visualization purposes. Bottom: Calcium changes in 9Aβ neurons during tibia vibration across different frequencies. Thin lines are calcium signals from individual flies, thicker line indicates the average across flies.

To ask whether 9Aβ neurons integrate club signals from multiple legs, we compared calcium responses from 9Aβ neurons in a single neuromere (ipsilateral or contralateral) while stimulating club axons from the left prothoracic leg using targeted optogenetic excitation. 9Aβ neurons in both the left and right neuromeres increased their calcium activity in response to excitation of club axons (Figure 2C). These data suggest that 9Aβ neurons integrate ipsilateral and contralateral signals from club neurons (Figure 2E). EM reconstruction of a 9Aβ neuron with a cell body in the opposite neuromere confirmed the existence of direct synaptic input from contralateral club axons (Figure 2D).

To understand how 9Aβ neurons encode leg movements *in vivo*, we recorded 9Aβ calcium signals while manipulating the femur-tibia joint of the fly’s left leg with a magnetic control system (Mamiya et al., 2018; Figure 2F). We observed phasic calcium signals of 9Aβ neurons in both ipsilateral and contralateral neuromeres, in response to tibia flexion and extension (Figure 2G). Similar to what we observed in the club Split-Gal4 line (Figure S1D), 9Aβ also exhibited higher baseline activity when the tibia was held at full extension compared to flexion; inspection of high-speed video suggests that this response is caused by vibrations produced by the fly’s resistance to passive tibia extension. Consistent with this hypothesis, 9Aβ neurons responded strongly to low-amplitude (0.1µm) vibration of the tibia (Figure 2H) at frequencies from 200-2000 Hz, similar to the club neuron population (Mamiya et al., 2018). Thus, 9Aβ neurons encode high frequency, low amplitude movement of the tibia, consistent with a role for sensing external substrate vibrations.

In summary, GABAergic 9Aβ neurons integrate club signals from left and right legs in the same segment to encode tibia movement and high frequency vibration (Figure 2E). 9Aβ neurons are thus positioned to provide inhibition to other neurons in the vibration processing pathway, or to mediate interactions between movement and vibration pathways.

### 10Bα, a VNC cell class downstream of the club, integrates bidirectional movements and vibration signaling from different legs across segments

We next switched our attention to 10Bα, the second candidate cell class whose anatomy suggests that it integrates club signals from multiple legs. Single 10Bα neurons with a cell body in T1 arborize within one neuromere, then cross the midline and arborize in the contralateral T2 neuromere (Figure 3A).

**Figure 3.**
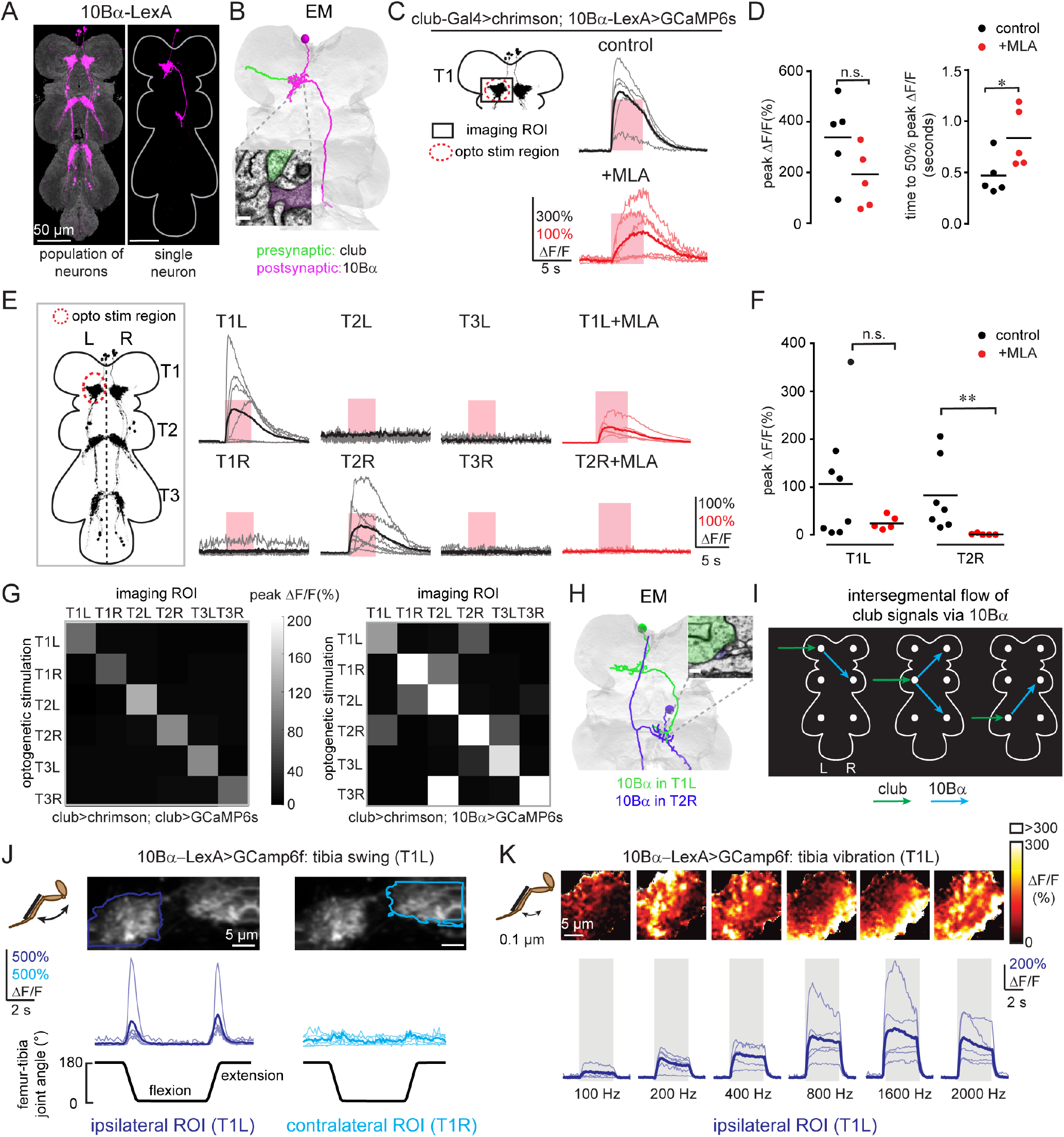
10Bα neurons integrate movement and vibration signals from club neurons across legs. (**A**) Anatomy of 10Bα neurons. Magenta is GFP driven by 10Bα (*R13E04-LexA*), while neuropil was stained by nc82. At right is a single neuron labeled by multi-color FLPout. Both images were aligned to a common VNC template. **(B)** Anatomical reconstruction from EM showing an example of a 10Bα neuron (magenta) that receives direct synaptic input from a club axon (green). The inset shows an example of a synapse between the two cells. Scale bar = 200nm. (**C**) Calcium responses of the 10Bα neurons to optogenetic stimulation of club neurons. Calcium responses of 10Bα neurons in the left prothoracic VNC (T1L) to stimulation of club neurons in the left foreleg. Methyllycaconitine (MLA, 1μM) effectively blocked excitation from club neurons. The pink windows indicate stimulus duration (5 seconds, laser power= 0.28 mW/mm_2_). (**D**) Peak calcium responses (left) and time to 50% of the maximum calcium signals (right) across flies, for the experiments shown in (C). Each dot represents data from a single fly, while bars represent average peak calcium signals (left) or mean time to peak (right) (control: n=5, MLA: n=5. * p<0.05, n.s. no significant difference, Mann-Whitney test). (**E**) Calcium (T1L-T3R) to stimulation of club axons in T1L with or without MLA (1µM). The pink windows indicate stimulus duration (5 seconds, laser power= 0.28 mW/mm^2^). (**F**) Same as in D but showing the quantification of the peak calcium responses shown in (E). Each dot represents data from a single fly (T1L: n=7, T2R: n=5, n.s.: no significance, * p<0.05, Mann-Whitney test). (**G**) Heatmaps of average peak calcium responses of club (left, n=5 flies) and 10Bα (right, n=6 flies) neurons in each neuropil to stimulation of the axons of club neurons in each leg. (**H**) Same as in B but showing a 10Bα neuron in T1 left (green) synapses on a 10Bα neuron in T2 right (blue) via EM reconstruction. Scale bar = 200nm. (**I**) Proposed diagram of signal flow from club axons to 10Bα neurons, based on data summarized in (G). White dots represent neurites of the 10Bα neurons in different neuromeres. (**J**) Calcium response in 10Bα neurons during tibia swing movement. 10Bα neurons respond phasically to bidirectional tibia movement (n=6). (**K**) 10Bα neurons respond to tibia vibration. Top: The majority of pixels had a ΔF/F value between 0 and 300%; outlier pixels with a value above 300% ΔF/F were set to white for visualization purposes. Bottom: Calcium changes in 10Bα neurons during tibia vibration across different frequencies. Thin lines are calcium signals from individual flies, while the thicker line indicates the average across flies (n=5).

A subset of 10Bα neurons also project to the brain, arborizing in the contralateral antennal mechanosensory and motor center (AMMC, data not shown). Previous work showed that optogenetic activation of 10Bα neurons caused walking flies to pause, consistent with a role in detecting substrate vibration (Agrawal et al., 2020).

We reconstructed 10Bα neurons in the EM volume and found dense synaptic inputs from club axons (Figure 3B), confirming that club neurons are both functionally and anatomically presynaptic to 10Bα neurons. However, blocking acetylcholine receptors with MLA reduced but did not eliminate club-driven calcium signals in 10Bα neurons (Figure 3C-D). These data suggest that the connection between club and 10Bα neurons consists of mixed chemical/electrical signaling, which is consistent with previous work (Agrawal et al., 2020). Interestingly, we observed that the time to peak of the 10Bα calcium signals was significantly longer in the presence of MLA (Figure 3D), which suggests that chemical and electrical transmission may have distinct temporal dynamics.

The intersegmental projections of 10Bα neurons raise the possibility that these cells integrate signals from club neurons across different legs. To test this, we measured calcium responses of 10Bα neurons in each neuromere (T1L-T3R) while optogenetically stimulating club axons arising from each of the six leg nerves. When club axons from the left front (T1L) leg were optogenetically stimulated, we observed robust calcium signals in 10Bα processes of T1L and T2R, but not in other neuromeres (Figure 3E). Applying MLA abolished calcium responses in T2R, but not in T1L (Figure 3E-F), suggesting a role for gap junctions in local but not intersegmental connectivity.

To test whether this connectivity pattern generalized to other VNC segments, we consecutively stimulated club axons from each leg while recording calcium signals from 10Bα neurons in all six neuromeres, resulting in a 6×6 functional connectivity map (Figure 3G). We observed intersegmental responses for all legs, though the pattern of information flow was different for each segment (Figure 3G, right). In contrast, stimulating and recording from the same club axons did not produce intersegmental responses (Figure 3G, left). These data demonstrate that 10Bα neurons integrate club signals from pairs of adjacent legs. This conclusion is also supported by our finding from EM reconstruction that 10Bα neurons from T1L and T2R are synaptically connected (Figure 3H).

To compare these functional connectivity results to encoding of sensory stimuli, we recorded calcium signals in 10Bα neurons while moving the tibia at 180°/sec. Unlike 9Aβ, 10Bα neurons responded to ipsilateral but not contralateral tibia movement in the prothoracic segment (Figure 3J). Like club axons (Figure S1D), 10Bα neurons responded to tibia movement in both directions (Figure 3J), as well as high frequency, low amplitude vibration of the tibia (Figure 3K). The distribution of calcium signals shifted from lateral to medial as vibration frequency increased (Figure 3K), consistent with the topographic map of frequency previously observed in club axons (Mamiya et al., 2018). Curiously, this frequency map was not present in recordings from 9Aβ neurons (Figure 2H).

In summary, both 9Aβ and 10Bα neurons encode bidirectional movements and vibration signals by integrating club axons across multiple legs. The key difference between these cell classes is that 9Aβ neurons mediate bilateral inhibition within a VNC segment, while 10Bα neurons integrate excitatory club inputs across contralateral VNC segments (Figure 3I). This convergence supports our hypothesis that club pathways play a role in detecting substrate vibration: external vibration stimuli are likely to be correlated across legs, in contrast to natural joint kinematics, which are unlikely to be correlated across legs.

### 13Bβ neurons integrate position and directional movement signal

Another interesting result of our functional connectivity screen was that some second-order neurons integrate proprioceptive signals across multiple FeCO subtypes. Specifically, we identified two candidate cell-types (13Bβ and 8Aα) downstream of both claw and hook axons. We selected 13Bβ neurons for further analysis because clean genetic driver lines exist for this cell class.

We reconstructed the anatomy of 13Bβ cells from the EM volume and found direct synaptic inputs from hook-extension axons (Figure 4B). We did not find any synapses between 13Bβ and claw neurons, probably because only 2 claw axons have been fully reconstructed; however, we cannot rule out the possibility that claw axons are connected to 13Bβ neurons indirectly. Calcium responses in 13Bβ neurons were abolished when we blocked acetylcholine receptors with MLA (Figure 4C), suggesting that the synaptic input from both FeCO subtypes relies on chemical synaptic transmission.

**Figure 4.**
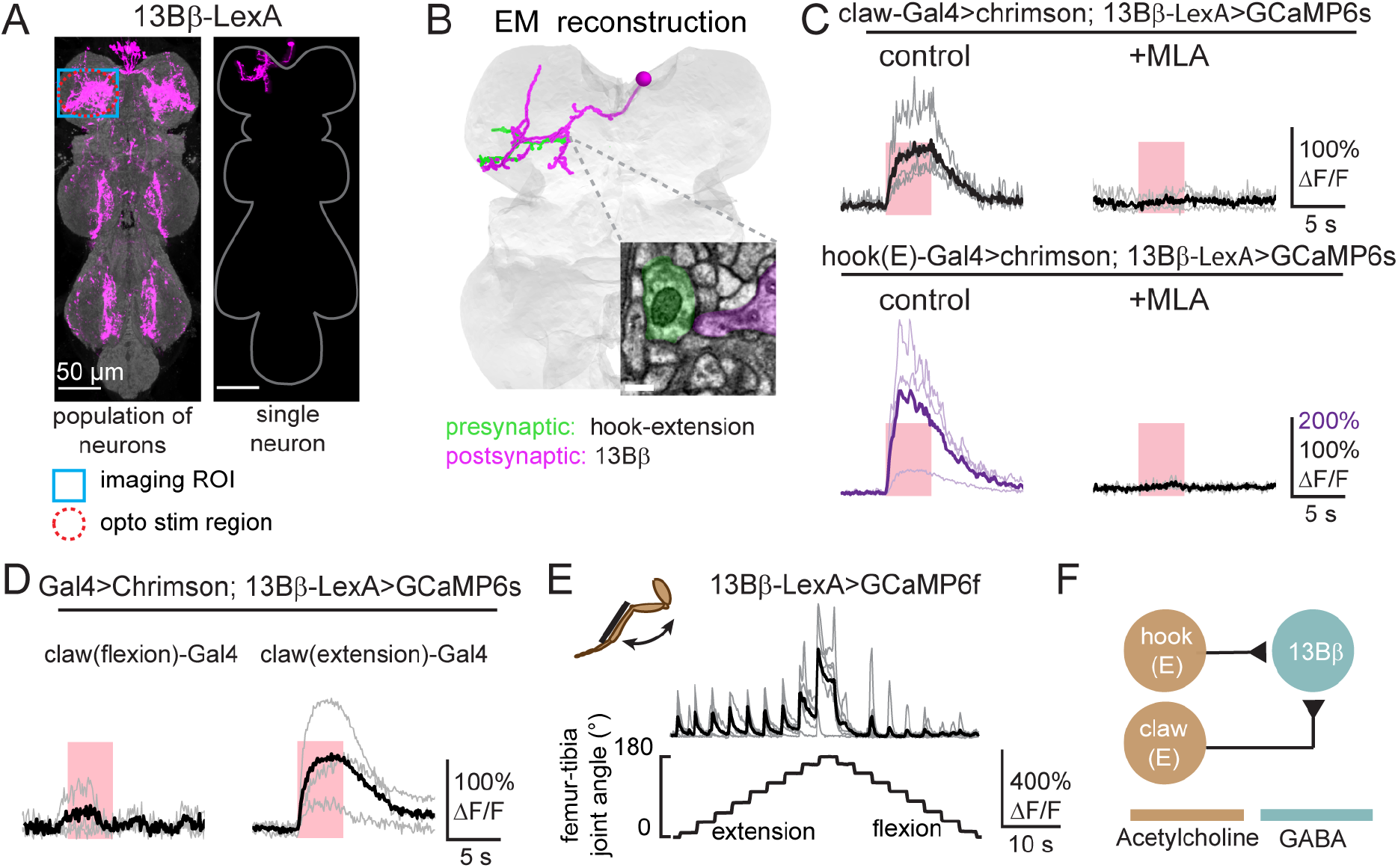
13Bβ neurons integrate position and directional movement signals from claw and hook-extension neurons. (**A**) Anatomy of 13Bβ neurons. Population (left) and single neuron (right) anatomy of 13Bβ neurons. GFP (magenta) was driven by 13Bβ (*VT006903-LexA*). The VNC neuropil was stained against nc82 (grey). Both images were aligned to a common VNC template. (**B**) Anatomical reconstruction using EM showing an example of a 13Bβ neuron (magenta) that receives direct synaptic input from a hook-extension axon (green). The inset shows an example of a synapse between the two cells. Scale bar is 200 nm. (**C**) Calcium responses of 13Bβ neurons to optogenetic stimulation of claw and hook-extension neurons. Left: calcium responses of 13Bβ neurons in the left prothoracic VNC to optogenetic stimulation of claw axons from the left foreleg (T1L). Right: MLA (1µM) blocked excitation produced by claw neuron activation. The pink windows indicate stimulus duration (5 seconds, laser power= 0.28 mW/mm^2^). Control: n=5, MLA: n=4 respectively. Bottom: calcium responses of 13Bβ neurons to optogenetic stimulation of hook-extension axons. (**D**) Calcium responses of 13Bβ to optogenetic stimulation of claw-flexion (n=3) and claw-extension axons (n=4). (**E**) 13Bβ neurons respond to tibia extension. Calcium changes of 13Bβ neurons during tibia movement (n=6). (**F**) Proposed diagram of sensory integration by 13Bβ neurons, which receive input from claw-extension and hook-extension neurons.

We next sought to understand the convergence of position and movement signals within 13Bβ neurons. Our functional connectivity screen revealed that 13Bβ neurons respond to activation of claw axons, but the driver line we used to activate these neurons labeled both flexion and extension-tuned cells. We therefore created Split-Gal4 lines that separately label claw neurons encoding tibia flexion (<90°) and extension (>90°; Figure S4) and repeated functional connectivity experiments with these sparser lines. 13Bβ neurons specifically increased calcium activity in response to optogenetic stimulation of extension-tuned claw neurons but did not respond to flexion-tuned claw neurons (Figure 4D). Calcium responses during passive tibia movements were consistent with convergent input from extension-tuned claw and hook neurons: 13Bβ calcium signals peaked during extension movements when the tibia was already extended (Figure 4E).

In summary, GABAergic 13Bβ neurons integrate excitatory input from extension-tuned claw and hook neurons to encode joint movement within a specific angular range (Figure 4F). Integration of direction and position signals could be beneficial for dynamic tuning of resistance reflexes that maintain body posture and protect joints from hyperextension.

### Inhibition gates calcium dynamics of central proprioceptive pathways

Our screen (Figure 1) identified several cell classes with dendrites in close proximity to FeCO axons, but which did not respond to optogenetic activation of proprioceptors. We wondered if their connectivity was masked by feedforward inhibition, as has been demonstrated in second-order neurons that process tactile signals from the leg (Tuthill and Wilson, 2016a). We repeated functional connectivity experiments with these cell classes while blocking GABA_a_ receptors with picrotoxin (10µM). From 8 VNC cell classes we tested, two (9Aγ and 20/22Aβ) had measurable calcium signals only in the presence of picrotoxin (Figure 5A). These connections were specific, meaning that 9Aγ neurons responded only to activation of club neurons and 20/22Aβ neurons responded only to activation of claw neurons.

**Figure 5.**
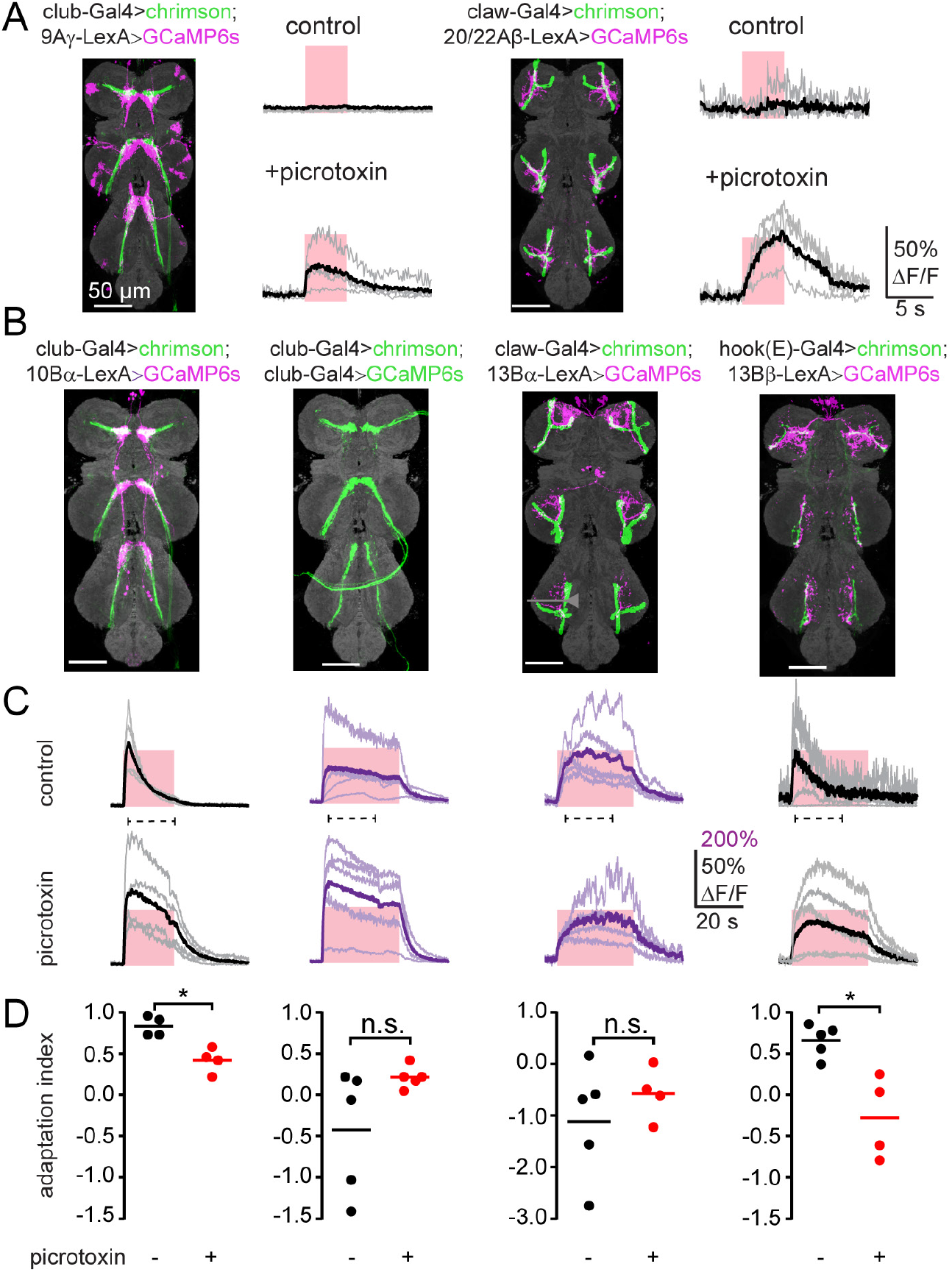
Multiple roles for inhibition in functional connectivity between first and second-order proprioceptive neurons. (**A**) Inhibition gates connectivity between leg proprioceptors and VNC neurons. Left: responses of 9Aγ neurons to optogenetic stimulation of club neurons were revealed after application of picrotoxin (10 µM). Right: similar results for claw and 20/22Aβ neurons. The pink windows indicate stimulus duration (5 seconds, laser power= 0.28 mW/mm^2^). (**B**) Anatomy of the axons of FeCO subtypes (green) and their downstream targets (magenta). VNC neuropil was stained using nc82 (grey). (**C**) Calcium responses of second-order neurons in the left prothoracic VNC to optogenetic stimulation of the indicated sensory neurons (top). Picrotoxin (10 µM) reduced response adaptation in 10Bα and 13Bβ neurons. The pink windows indicate the optogenetic stimulus duration (20 sec for 10Bα neurons and 30 sec for others; the laser power was 0.28mW/mm^2^). The dashed line under each trace indicates the window used to calculate the adaptation index below. (**D**) Quantification of calcium signal adaptation from data in (C). Adaptation index was calculated as 1-F_offset_/F_peak_. 1 indicates complete adaptation, 0 indicates no adaptation, negative values indicate an increase of the calcium signal over time. Each dot represents data from a single fly, while bars represent the averages (* p<0.05, n.s. no significant difference, Mann-Whitney test.).

In other cell classes, we found that GABAergic inhibition sculpted the dynamics of the calcium response. For example, GCaMP signals recorded from 10Bα neurons adapted during prolonged optogenetic stimulation (20 seconds) of club neurons (Figure 5C). In contrast, GCaMP signals in 13Bα neurons remained elevated during optogenetic stimulation of claw neurons over a period of 30 seconds (Figure 5C). This adaptation was not due to decay of optogenetically-evoked activity in the proprioceptor axons. Rather, it appears that inhibition, likely mediated by GABA_a_ or GluCl receptors (Liu and Wilson, 2013), contributes to the adaptation of 10Bα calcium signals during prolonged stimulation. We observed a similar phenomenon for 13Bβ neurons during optogenetic stimulation of hook-extension neurons (Figure 5C). Overall, these results show that adaptation mediated by GABAergic or glutamatergic inhibition gates the activity and sculpts the dynamics of second-order proprioceptive neurons.

## Discussion

In this study, we report the anatomical structure and functional organization of second-order circuits for leg proprioception in *Drosophila*. Due to the lack of clear hierarchical structure within the VNC leg neuropil, it has been challenging to infer the flow of proprioceptive sensory signals with existing tools. We therefore generated genetic driver lines to label specific subtypes of leg proprioceptors and classified candidate second-order neurons based on hemilineage identity. We used optogenetics and calcium imaging to map the functional connectivity between leg proprioceptors and second-order neurons, followed by EM reconstruction to validate synaptic connectivity. We then used spatially targeted and subtype-specific optogenetic stimulation to analyze integration of FeCO signals within a subset of second-order neuron classes.

Overall, this work reveals the logic of sensory integration in second-order proprioceptive circuits: some populations of second-order neurons integrate tibia vibration signals across pairs of legs, suggesting a role for detection of external substrate vibration. Signals for leg joint position and directional movement converge in other second-order neurons, revealing pathways for local feedback control of leg posture. We anticipate that this functional wiring diagram (Figure 6) will also help guide the interpretation of anatomical wiring diagrams determined through EM reconstruction of VNC circuits.

**Figure 6.**
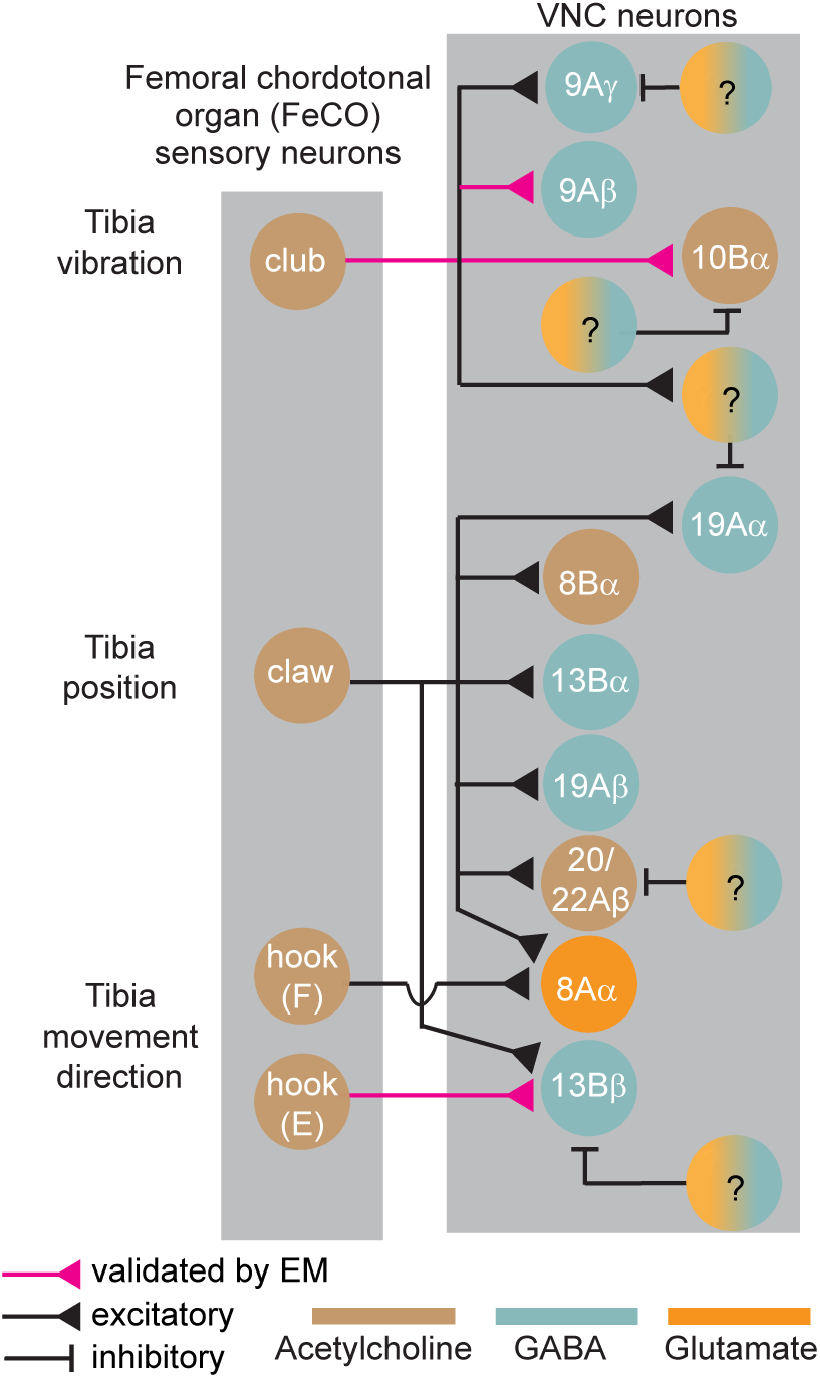
Summary diagram of circuits processing leg proprioceptive signals from the *Drosophila* FeCO, based on experiments in this study. Question marks indicate putative inhibitory neurons of unknown identity.

### Connectivity motifs within second-order proprioceptive circuits

Proprioceptors in the *Drosophila* FeCO can be classified into three subtypes: club neurons encode bidirectional tibia movement and vibration frequency, claw neurons encode tibia position (flexion or extension), and hook neurons encode the direction of tibia movement (Mamiya et al., 2018). Our results show the existence of two distinct central pathways for processing signals from club and claw/hook neurons (Figure 1). We propose that neurons downstream of the club mediate sensing of low-amplitude mechanical vibrations in the external environment, while neurons downstream of the claw and hook provide proprioceptive feedback to motor circuits for controlling the posture and movement of the legs. This division of central pathways for external and internal sensing may be a common motif across limbed animals. Work in a variety of species, including a recent study in mice (Prsa et al., 2019), has found that many animals can detect low-amplitude, high-frequency substrate-borne vibrations (Hill and Wessel, 2016). Flies may use vibration sensing to monitor acoustic signals in the environment, such as during courtship behavior (Fabre et al., 2012), or to detect approaching threats.

The distinct anatomical organization of neurons downstream of the club and claw/hook also supports a segregation of vibration sensing and motor control feedback pathways. 9Aβ and 10Bα neurons arborize in the medial ventral association center (mVAC, Figure 1E), a common target of descending neuron axons (Namiki et al., 2018). In contrast, 13Bβ arborize in the intermediate neuropil (IntNp, Figure 1E), which contains the dendritic branches of the leg motor neurons and premotor interneurons. Based on these differences, we hypothesize that vibration-sensing neurons interact with ascending and descending signals to/from the brain, while neurons downstream of hook and claw axons contribute to local motor control through direct or indirect connections to motor neurons. Leg motor neurons receive position and movement-tuned proprioceptive input, consistent with feedback from claw and hook neurons (Azevedo et al., 2020). Additional connectomic reconstruction is needed to determine which second-order neurons mediate these feedback connections, but 13Bβ neuros are promising pre-motor candidates.

We found that VNC neurons postsynaptic to claw and hook axons receive only local input, from individual legs, while second-order neurons postsynaptic to club axons integrate signals across multiple legs. For example, GABAergic 9Aβ neurons pool information from left and right legs in a single VNC segment (Figure 2), while cholinergic 10Bα neurons receive convergent input from left and right legs across different segments (Figure 3). Integrating club input across legs may improve detection of external vibration signals, while proprioceptive signals from the claw and hook may be initially processed in parallel to support reflexive motor control of individual legs. Bilateral integration also occurs in second-order auditory circuits downstream of the *Drosophila* Johnston’s organ: mechanosensory signals from the two antennae are processed in parallel by second-order neurons in the AMMC, but then converge in third-order neurons in the wedge (Patella and Wilson, 2018).

While second-order neurons in the vibration pathway integrate club signals across legs, multiple classes of second-order neurons in the motor pathway (13Bβ and 8Aα) integrate signals across different FeCO subtypes from the same leg. Using new genetic driver lines that subdivide claw neurons into extension and flexion-tuned subtypes, we found that extension-tuned claw and hook neurons converge on 13Bβ neurons. We hypothesize that these cells mediate resistance reflexes that stabilize tibia position in response to external perturbations. Prior work in the stick insect has shown that tibia resistance reflexes rely on position and directional movement signals from the FeCO (Bässler, 1993). In support of this hypothesis, another class of neurons in the 13B hemilineage, 13Bα, also encode tibia extension and drive tibia flexion when optogenetically activated (Agrawal et al., 2020).

### Inhibition and temporal dynamics

Our results identify a prominent role for inhibition in central processing of proprioceptive information from the FeCO. Of the eight second-order cell classes we identified in our screen, six are putative inhibitory neurons (i.e., release GABA or glutamate). In other sensory circuits, local inhibitory processing contributes to sharpening spatial and temporal dynamics (Dubs et al., 1981) as well as reducing sensory noise through crossover inhibition (Cafaro and Rieke, 2013; Liu et al., 2015). By pharmacologically blocking GABA_a_ and GluCl receptors, we identified a role for inhibition in controlling adaptation within second-order neurons (e.g., 10Bα and 13Bβ neurons, Figure 5C). In other cases (20/22Aα or 9Aγ, Figure 5A), inhibition was strong enough to completely mask proprioceptive inputs from FeCO axons. We hypothesize that this inhibition may be tuned in certain behavioral contexts, for example during active movements, to gate the flow of proprioceptive feedback signals in a context-dependent manner.

Synaptic transmission in *Drosophila* can be mediated by chemical synapses, which can be visualized with EM, or electrical gap junctions, which are not typically identifiable at the resolution of current EM volumes. FeCO neurons release acetylcholine but also express gap junction proteins (*shakB*, data not shown). We therefore used pharmacology to test for the presence of gap junctions between sensory and central neurons. MLA, an effective antagonist of nicotinic acetylcholine receptors in *Drosophila* (Tuthill and Wilson, 2016a), eliminated functional connectivity between club and 9Aβ neurons, but only reduced functional connectivity between club and 10Bα neurons. We observed similar results downstream of the claw: MLA blocked functional connectivity between claw and 13Bβ neurons, but only reduced functional connectivity between claw and 13Bα neurons (data not shown). These results suggest that second-order proprioceptive circuits receive mixed chemical and electrical input from FeCO neurons. More work is needed to confirm these observations and to investigate the functional significance of why pathways might use one means of signal transmission over the other. One hypothesis is that chemical synapses exhibit adaptation (e.g., synaptic depression) while electrical synapses may be more advantageous for sustained synaptic transmission (Grimes et al., 2014). Each may provide different advantages for pathways that control behavior on a variety of timescales, from slow postural reflexes to rapid escape behaviors.

### Comparison to central mechanosensory processing in other species

Central processing of sensory signals from the FeCO has been previously studied in other insects, especially the locust (Burrows, 1996) and stick insect (Bässler and Büschges, 1998). In both species, second-order interneurons encode combinations of tibia movement and position (Buschges, 1989), and also integrate multimodal signals from different proprioceptive organs (Burrows, 1985; Siegler and Burrows, 1983). Vibration signals detected by the FeCO appear to be processed by largely segregated populations of VNC interneurons (Büschges, 1989; Stein and Sauer, 1999). However, these conclusions were based on mapping of sensory receptive fields, and it was not previously possible to identify specific sources of sensory input, as we do in this study.

Overall, comparison of our functional connectivity results in *Drosophila* with the prior work in other insect species suggests broad evolutionary conservation of VNC circuits for leg proprioception and motor control. Although it is currently difficult to identify homologous cell-types across insect species, future efforts could leverage conserved developmental programs: the organization of neuroblasts that give rise to the VNC is similar across insect species separated by 350 million years of evolution (Lacin and Truman, 2016). This is an important advantage of using developmental lineages to define VNC cell classes – locusts and stick insects also possess 9A, 10B, and 13B neurons, which could someday be identified based on molecular markers of lineage identity.

### Complementary strengths of functional and structural connectivity

The functional connectivity approach that we employed in this study has both benefits and drawbacks. On the positive side, it allowed us to screen a large connectivity matrix of genetically-identified sensory and central neurons. Compared to other methods for anatomical mapping (e.g., EM), the use of optogenetics and calcium imaging allowed us to measure connection strength and dynamics across multiple individuals. We found that second-order VNC neurons varied significantly in their functional connectivity strength and temporal dynamics (Figure 1G-H). We also observed 5-fold differences in peak calcium signals in response to optogenetic stimulation with the same light intensity (Figure 1G). This range could be due to differences in GCaMP expression or intracellular calcium buffering but may also reflect differences in synaptic strength between pre-and postsynaptic partners.

On the other hand, the functional connectivity method cannot resolve whether inputs are direct, due to the slow kinetics of GCaMP6. We therefore used sparse, targeted EM tracing to validate some of the functional connections we identified between FeCO and VNC neurons. A more detailed comparison of functional and anatomical connectivity will require dense, comprehensive reconstruction of the VNC neuropil. Automated reconstruction and manual proofreading have recently led to draft wiring diagrams of neural circuits in the adult *Drosophila* central brain (Scheffer et al., 2020). Similar approaches to reconstruct the VNC connectome are in progress (Phelps et al., 2021).

## Acknowledgements

We thank Haluk Lacin and members of the Tuthill lab for comments on the manuscript. We thank Haluk Lacin, David Shepherd, and Jim Truman with assistance identifying VNC lineages. We thank Allan Wong and Daniel Bushey for help with 2-photon microscopy, the Janelia Fly Facility, and the Project Technical Resources for technical assistance. Stocks obtained from the Bloomington *Drosophila* Stock Center (NIH P40OD018537) were used in this study. This work was supported by the Visiting Scientist Program at HHMI/Janelia, a Searle Scholar Award, a Klingenstein-Simons Fellowship, a Pew Biomedical Scholar Award, a Sloan Research Fellowship, the New York Stem Cell Foundation, and NIH grant R01NS102333 to JCT. JCT is a New York Stem Cell Foundation – Robertson Investigator.

## Material and Methods

### Fly stocks

*Drosophila* were raised on cornmeal agar food on a 12h dark/12h light cycle at 25°C. Female flies, 4-8 days post eclosion, were used for all calcium imaging experiments. For functional connectivity experiments, adult flies carrying the Chrimson transgene were placed on cornmeal agar with all-trans-retinal (0.2 mM, dissolved in 95% EtOH, Sigma-Aldrich) for 2-3 days prior to the experiment. Vials were wrapped in aluminum foil to reduce unnecessary optogenetic activation.

### Creation of Split-GAL4 lines for targeting proprioceptors in fly leg

GAL4 images from the Rubin and Dickson collections (Jenett et al., 2012; Tirian and Dickson, 2017) were visually screened for lines labeling axons from proprioceptors that project to the VNC. For each cell type, a color depth MIP mask search (Otsuna et al., 2018) was performed to find other GAL4 lines with expression in similar cells. Split-GAL4 AD and DBD hemi-drivers (Dionne et al., 2018; Tirian and Dickson, 2017) for these lines were crossed in several different combinations to identify intersections that targeted the cell type of interest but with minimal expression elsewhere. Sensory tuning properties of FeCO subclass neurons labeled by these Split-Gal4 lines were further characterized using *in vivo* calcium imaging, described below.

### Immunohistochemistry and anatomy

For confocal imaging of the FeCO neuron axons driven by each Split-Gal4 line in the VNC (Figure S1, S4), we crossed flies carrying the Split-Gal4 driver to flies carrying 20xUAS-IVS-mCD8::GFP or 20xUAS-Chrimson::mVenus (attp18) and dissected the brain and VNC of the resulting progeny in cold Schneider’s Insect Medium (S2). The tissues were first fixed in 2% paraformaldehyde (PFA) PBS solution for 55 min followed by rinsing in PBS with 0.5% Triton X-100 (PBT) four times. The brain and VNC were blocked in solution (5% normal goat serum in PBT) for 90 min, then incubated with a solution of primary antibody (rabbit anti-GFP 1:1,000 concentration; mouse nc82 for neuropil staining; 1:30 concentration) in blocking solution for 4 hrs, followed by washing tissues in PBT three times. Tissues were incubated with a solution of secondary antibody (anti-rabbit-Alexa 488 1:400 concentration; anti-mouse-Alexa 633 1:800 concentration) dissolved in blocking solution for 4 hrs followed by three times washing with PBT before DPX mounting. The whole procedure was performed at room temperature.

For stochastic labeling of individual neurons in the VNC, we crossed flies carrying the multicolor FlpOut cassettes and Flp recombinase drivers to flies carrying different Split-Gal4 and LexA drivers and dissected out the VNCs of resulting progeny. For temperature induced expression of Flp, we placed adult flies at 1-3 days old in a plastic tube and incubated them in a 37°C water bath for 15 min. We dissected the VNC four days after the Flp induction and followed the procedure described in (Nern et al., 2015) to detect HA (using anti-HA-rabbit antibody and anti-Rabbit-Alexa 594 secondary antibody), V5 (using DyLight 549-conjugated anti-V5), and FLAG (using anti-FLAG-rat antibody and anti-Rat-Alexa 647 secondary antibody) labels expressed due to Flp induction in individual neurons.

Images were acquired on Zeiss LSM 710 or 800 confocal microscopies with 20x or 63x objectives. We used Fiji (Schindelin et al., 2012) to generate maximum intensity projections of the expression of driver lines as well as anatomy of single neurons.

### Fly preparation for two-photon calcium imaging

For functional connectivity experiments, adult female flies were anesthetized on ice and then glued to a petri dish with ventral side up using UV-cured glue (Kemxert 300). To eliminate spontaneous activity due to fly leg movement, we amputated the legs at the coxa joint. After immersing the fly in extracellular fly saline (103mM NaCl, 3mM KCl, 2mM CaCl_2_, 4mM MgCl_2_, 26mM NaHCO_3_, 1mM NaH_2_PO_4_, 8mM trehalose, 10mM glucose, 5mM TES, pH 7.1, osmolality adjusted to 270-275 mOsm, bubbled with 95% O_2_ / 5% CO_2_), we removed the cuticle above the prothoracic segment of the VNC and took out the digestive tract to reduce the movement of the VNC. Recordings were performed at room temperature.

For calcium imaging during controlled leg movements, we used a fly holder previously described by Mamiya et al. (2018). Flies were anesthetized on ice and then positioned ventral side up, with the head glued to the upper side of the fly holder using UV-cured glue (Kemxert 300). We glued the ventral side of the thorax onto the hole and on the bottom side of the holder and glued down the femur of the right prothoracic leg so that we could control the femur-tibia joint angle by moving the tibia. When gluing the femur, we held it at a position where the movement of the tibia during the rotation of the femur-tibia joint was parallel to the plane of the fly holder. To eliminate mechanical interference, we also glued down the other legs. We pushed the abdomen to the left side and glued it at that position, so that the abdomen did not block tibia flexion. To position the tibia using the magnetic control system described below, we cut a small piece of insect pin (length ∼1.0 mm, 0.1 mm diameter; Living Systems Instrumentation) and glued it onto the tibia and the tarsus of the right prothoracic leg. To enhance contrast and improve tracking of the tibia/pin position, we painted the pin with black India ink (Super Black, Speedball Art Products). After immersing the ventral side of the preparation in extracellular fly saline, we removed the cuticle above the prothoracic segment of the VNC and took out the digestive tract to reduce the movements of the VNC. We also removed fat bodies and larger trachea to improve access to the leg neuropil. Fly saline contained: 103mM NaCl, 3mM KCl, 2mM CaCl_2_, 4mM MgCl_2_, 26mM NaHCO_3_, 1mM NaH_2_PO_4_, 8mM trehalose, 10mM glucose, 5mM TES, pH 7.1, osmolality adjusted to 270-275 mOsm. Recordings were performed at room temperature.

### Image acquisition using a two-photon excitation microscope

For functional connectivity experiments, images were acquired using a two-photon microscope (custom-made at Janelia by Dan Flickinger and colleagues, with a Nikon Apo LWD 25× NA1.1 water immersion objective). The standard imaging mode was a 512 × 512 image at 2.5 frames/s, and a ∼353 μm × ∼353 μm field of view (∼0.69 μm × ∼0.69 μm / pixel). The sample was imaged using a near-infrared laser (920nm, Spectra Physics, Insight DeepSee) that produced minimal activation of Chrimson at our typical imaging power (4-10 mW). Chrimson was activated by 590nm light (Thorlabs M590L3-C1) presented through the objective. Photoactivation light was delivered in a pulse train that consisted of six 5s pulses (within each 5 s pulse: square-wave modulation at 50 Hz, 30s inter-pulse interval). The light intensity increased for each of the six pulses (0.02, 0.04, 0.12, 0.28, 0.37, 0.68 mW/mm^2^). For targeted stimulation (e.g., Figure 2C, 3E), illumination was spatially modulated using a DMD (Digital Micromirror Device, Texas Instruments, DLP LightCrafter v2.0), and restricted to a specific stimulation region.

For calcium imaging during controlled leg movements, we used a modified version of a custom two-photon microscope previously described in detail (Euler et al., 2009). For the excitation source, we used a mode-locked Ti/sapphire laser (Mira 900-F, Coherent) set at 930 nm and adjusted the laser power using a neutral density filter to keep the power at the back aperture of the objective (40x, 0.8 NA, 2.0 mm wd; Nikon Instruments) below ∼25 mW during the experiment. We controlled the galvo laser scanning mirrors and the image acquisition using ScanImage software (version 5.2) within MATLAB (MathWorks). To detect fluorescence, we used an ET510/80M (Chroma Technology Corporation) emission filter (GCaMP6f or GCaMP6s) and a 630 AF50/25R (Omega optical) emission filter (tdTomato) and GaAsP photomultiplier tubes (H7422P-40 modified version without cooling; Hamamatsu Photonics). We acquired images (256 × 120 pixels or 128 × 240 pixels) at 8.01 Hz. At the end of the experiment, we acquired a z-stack of the labelled neurons to confirm the recording location.

### Image processing and calculating ΔF/F

We performed all calcium image processing and analyses using scripts written in MATLAB (MathWorks). After acquiring the images for a trial, we first applied a Gaussian filter (size 5×5 pixel, Σ = 3) and intensity threshold to minimize background noise. For calculating the GCaMP6 fluorescence change relative to the baseline (ΔF/F), we used the lowest average fluorescence level in a 10-frame window as the baseline fluorescence during that trial. For cases in which calcium signals were reduced relative to baseline (e.g., 19Aα neurons), we used the average fluorescence level in a 10-frame window at the beginning of each trial as the baseline. Because not all flies co-expressed tdTomato, we did perform image registration to correct for sample movement. From those flies that did co-express tdTomato, we observed that movement of the VNC was negligible.

We defined three parameters to analyze the temporal dynamics of calcium signals, as shown in Figure 1G: peak ΔF/F during the stimulation window, the time after stimulation at which the ΔF/F reaches 50% of the peak value (Figure 1I), and the half-decay time after the peak ΔF/F is reached (Figure 1J). For quantification of adaptation in Figure 5D, we calculated an adaptation index as 1-F_offset_/F_peak_, where F_peak_ indicates the peak ΔF/F, and F_offset_ is ΔF/F 19 seconds after the stimulus onset (where the stimulation offset typically occurs in club/10Bα neurons). An adaptation index of 1 would indicate 100% decay to baseline, while an index of 0 would indicate no adaptation. Negative index values indicate an increase in the calcium signal over time.

### Moving the tibia using a magnetic control system

We used a previously described magnetic control system (Mamiya et al., 2018) to manipulate the femur/tibia joint angle. To move the tibia/pin to different positions, we attached a rare earth magnet (1 cm height x 5 mm diameter column) to a steel post (M3×20 mm flat head machine screw) and controlled its position using a programmable servo motor (SilverMax QCI-X23C-1; Max speed 533,333°/s, Max acceleration 83,333.33°/s^2^, Position resolution 0.045°; QuickSilver Controls). To move the magnet in a circular trajectory centered at the femur-tibia joint, we placed the motor on a micromanipulator (MP-285, Sutter Instruments) and adjusted its position while visually inspecting the movement of the magnet and the tibia using the tibia tracking camera described below. For each trial, we controlled the speed and the position of the servo motor using QuickControl software (QuickSilver Controls). During all trials, we tracked the tibia position (as described below) to confirm the tibia movement during each trial. Because it was difficult to fully flex the femur-tibia joint without the tibia/pin and the magnet colliding with the abdomen, we only flexed the joint up to ∼18°. We set the acceleration of the motor to 72,000°/s^2^ for all ramp and hold and swing movements. Movements of the tibia during each trial varied slightly due to differences in the length of the magnetic pin and the positioning of the tibia and motor.

### Tracking the femur-tibia joint angle during imaging experiments

To track the position of the tibia, we illuminated the tibia/pin with an 850 nm IR LED (M850F2, ThorLabs) and recorded video using an IR sensitive high-speed video camera (Basler Ace A800-510um, Basler AG) with a 1.0x InfiniStix lens (94 mm wd, Infinity). The camera was equipped with a 900 nm short pass filter (Edmund Optics) to filter out the two-photon laser light. To synchronize the tibia movement with the recorded cell activity, the camera exposure signal and the position of the galvo laser scanning mirrors were acquired at 20 kHz. To track the tibia angle, we identified the position of the painted tibia/pin against the contrasting background by thresholding the image. We then approximated the orientation of the leg as the long axis of an ellipse with the same normalized second central moments as the thresholded image.

### Vibrating the tibia using a piezoelectric crystal

To vibrate the tibia at high frequencies, we moved the magnet using a piezoelectric crystal (PA3JEW, Max displacement 1.8 μm; ThorLabs). To control the movement of the piezo, we generated sine waves of different frequencies in MATLAB (sampling frequency 10 kHz) and sent them to the piezo through a single channel open-loop piezo controller (Thorlabs). Piezo-induced tibia movements during the calcium imaging prep were calibrated as described by Mamiya et al. (2018). For each stimulus, we presented 4 s of vibration 2 times with an inter-stimulus interval of 8 s. We averaged the responses within each fly before averaging across flies.

### Pharmacology

Drugs were added to the bath with a micropipette. Picrotoxin (Sigma-Aldrich) was prepared as a concentrated stock solution in aqueous NaCl (140 mM), and methyllycaconitine citrate (MLA, Sigma-Aldrich) were prepared as stock solutions in water. Each drug was further diluted in saline for experiments for a final concentration of 1 μM (MLA), or 10 μM (picrotoxin). The VNC was incubated in the drug for 5 min, with the perfusion system off, before starting the experiment, which typically lasted ∼30 min.

### Statistical analysis

For functional connectivity results in Figures 1-6, no statistical tests were performed *a priori* to decide on sample sizes, but sample sizes are consistent with conventions in the field. We used the Mann-Whitney non-parametric test to test for differences between two groups (Figure 1H,1I 3D, 3F, 5D), and the Kruskal-Wallis non-parametric test to test for differences between more than two groups (Figure 1I, 1J). All statistical analysis was performed with GraphPad (Prism). Results of statistical tests are reported in the figure legends.

### EM reconstruction

A TEM volume of the adult female VNC (Phelps et al., 2021) was used for reconstruction of neurons and their synaptic connectivity. FeCO axons were traced manually using CATMAID, a collaborative manual tracing environment (Schneider-Mizell et al., 2016). Second-order neurons were identified and matched to light-level data based on common hemilineage characteristics: primary neurite fasiculation, dendritic arborizations, and axon projections (Truman et al., 2010). Neurons were segmented in EM (methods in preparation), proofread in Neuroglancer (https://github.com/google/neuroglancer), skeletonized, and imported to CATMAID, where further proofreading was conducted. Postsynaptic sites on VNC neurons were identified by the presence of a dark post-synaptic density and a corresponding T-bar on the presynaptic cell, consistent with standards in the field (Zheng et al., 2018). Beginning with a synapse on each VNC neuron, sensory neurons were traced from the synapse back to the incoming axon such that they could be identified. Tracing was conducted until at least 3 synapses were found between each pair of neurons. We focused on identifying a minimal basis for connectivity between first and second-order neurons, due to ongoing efforts to automatically segment the entire VNC volume. In summary, we identified 3 synapses between a pair of club and 9Aβ neurons (both ispi- and contralateral connections), 3 synapses between a pair of club and 10Bα neurons, 3 synapses between a pair of 10Bα neurons in T1L and T2R, and 5 synapses between a pair of hook-extension and 13Bβ neurons.

## Data and software availability

Data and analysis code will be made available from the <u>authors website</u>.

## Supplemental Figures

**Figure S1.**
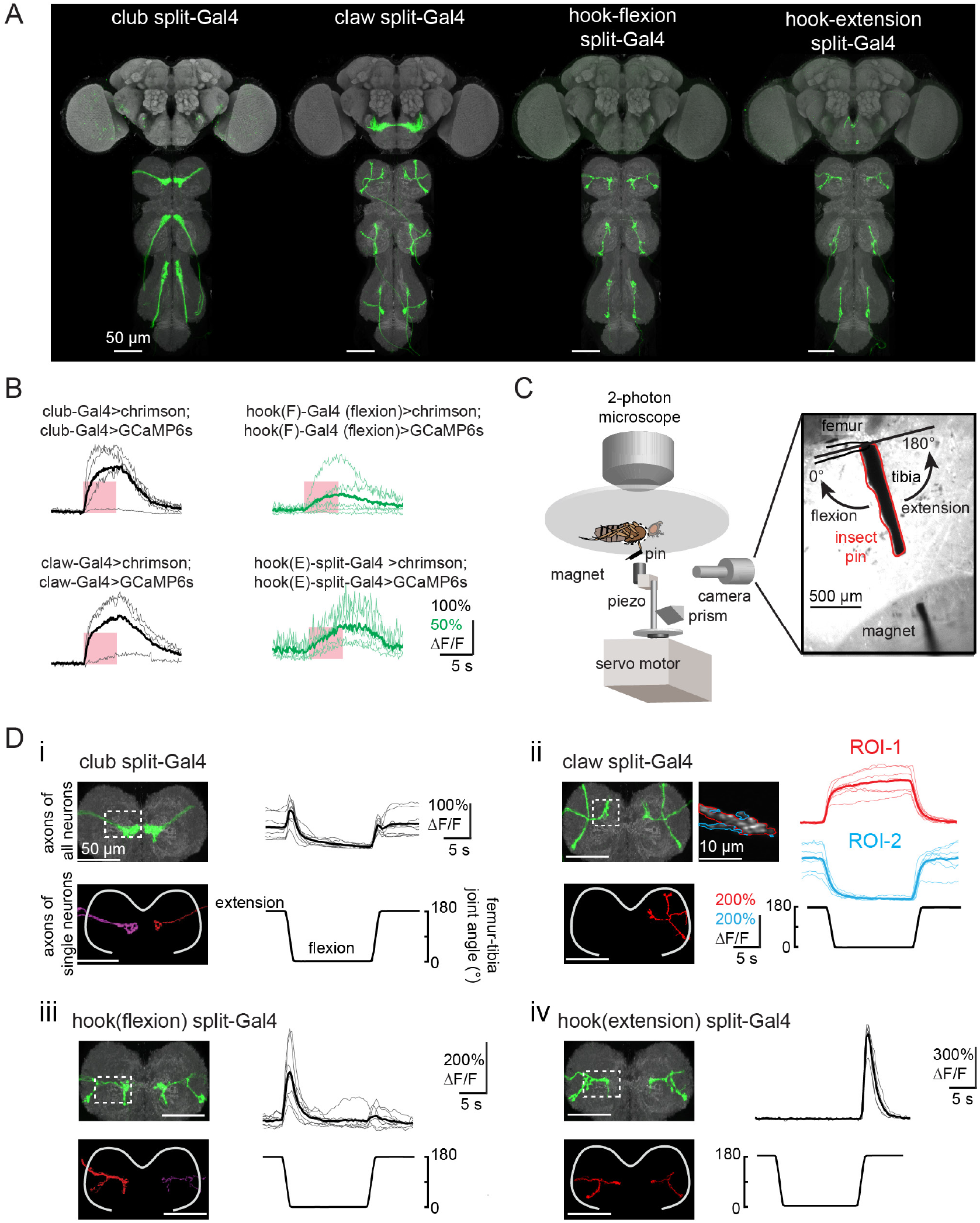
Split-Gal4 lines for targeting subtypes of femoral chordotonal organ (FeCO) proprioceptors. (**A**) GFP (green) expression in VNC and brain driven by split-Gal4 lines targeting subtypes of FeCO neurons. Grey: neuropil of VNCs and brains were stained with nc82. (**B**) Optogenetic stimulation of FeCO axons increases their calcium activity. Changes of GCaMP6s fluorescence relative to baseline (ΔF/F) in the axons of each FeCO subtypes to their self-stimulation. The thick lines in each panel represent average values. The pink windows indicate stimulus duration (5 seconds, laser power= 0.28 mW/mm^2^). (**C**) Experimental set-up for recording proprioceptive tuning of FeCO axons (adapted from Mamiya et al., 2018). (**D**) Anatomy and proprioceptive tuning of FeCO neurons labeled by four split-Gal4 lines. (i) Left: GFP labelled populations (upper panel) and single axon (lower panel) of club axons labeled by a split-Gal4 line. Grey: neuropil stained with nc82. Right: club neurons respond to bidirectional movement phasically. Tonic response at 180°caused by active tibia vibration at tibia fully extension. Changes of GCaMP7f fluorescence relative to baseline (ΔF/F) recorded from the regions outlined in a white rectangle at left when swung the tibia at 360°/s (n=8). (ii) Same as i, but for claw neurons responding tonically to tibia movement (n=7). Two sub branches could be further separated in response to flexion (ROI-1) and extension (ROI2) (iii) Same as (i), but for hook neurons that respond phasically to tibia flexion (n=7). (iv) Same as (i), but for hook extension neurons that phasically respond to tibia extension (n=6).

**Figure S2.**
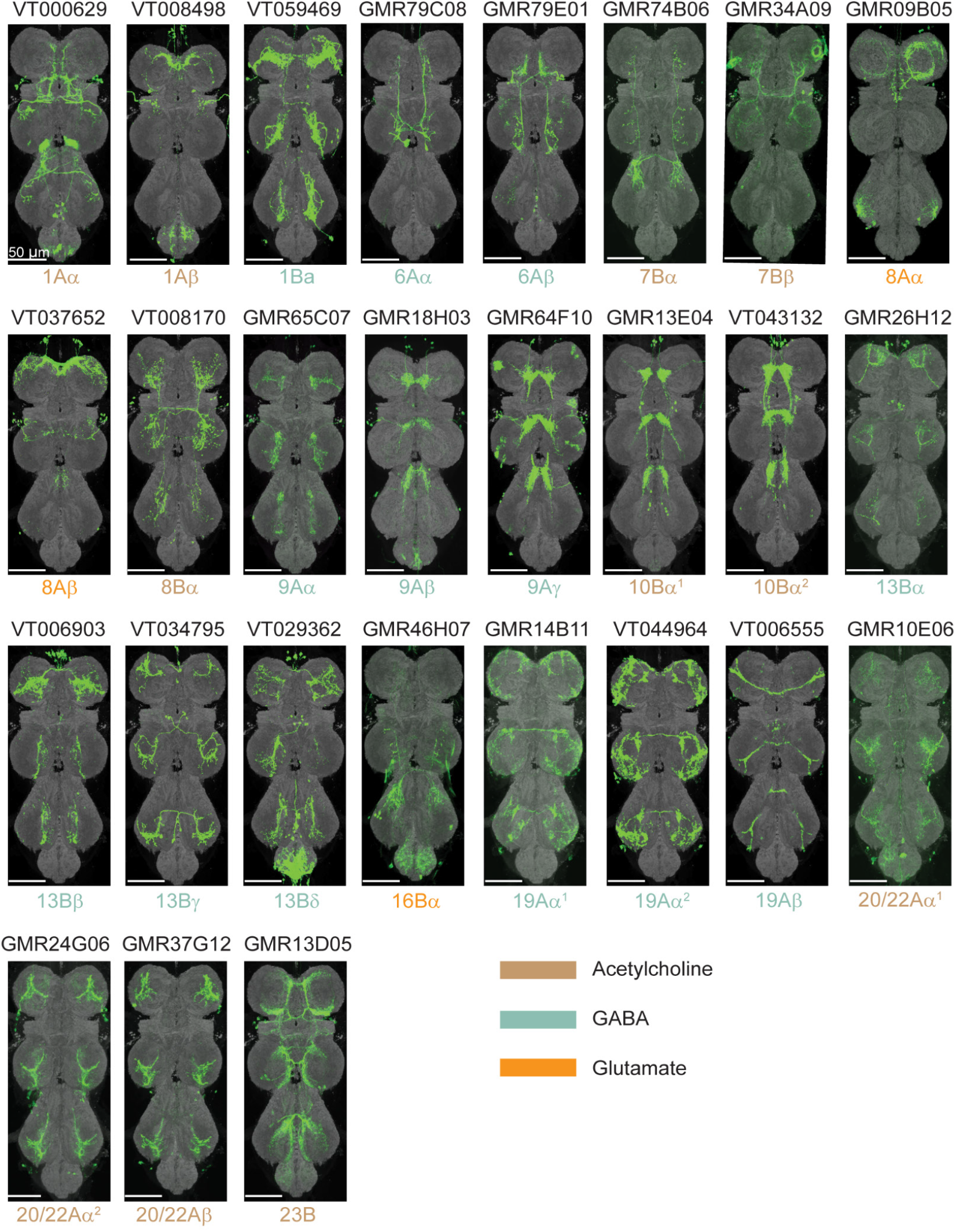
A collection of LexA driver lines used for functional connectivity experiments in this study. GFP (green) was expressed in the VNC driven by indicated LexA lines. Anatomy was used to determine the lineage identity described below each VNC image. The colors for each lineage and FeCO subtype indicate the putative neurotransmitter that they release.

**Figure S3.**
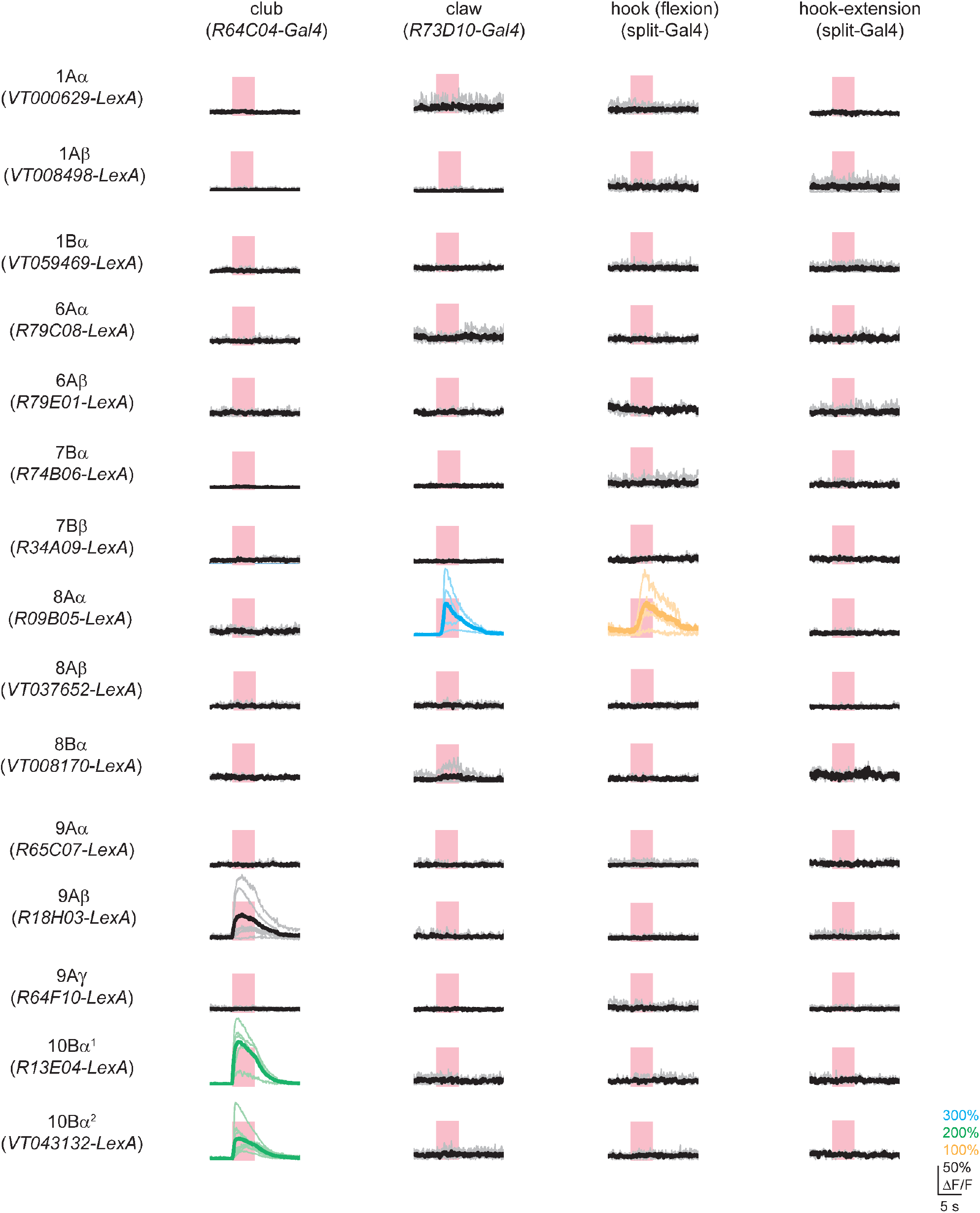
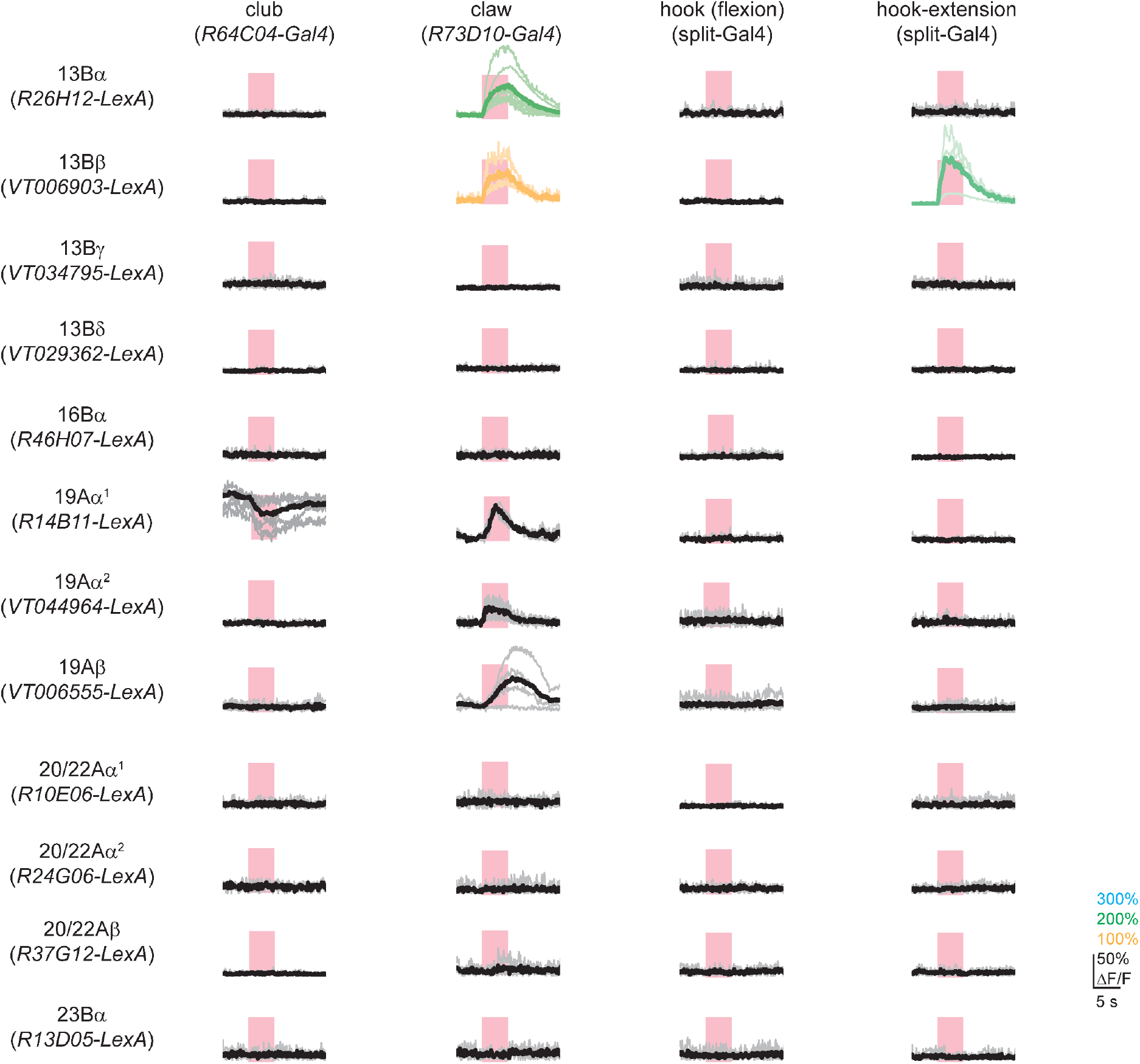
Times series data from functional connectivity experiments. Changes of GCaMP6s fluorescence relative to baseline (ΔF/F) were recorded in each driver line in response to optogenetic stimulation of four FeCO subtypes. The pink windows indicate stimulus duration (5 seconds, laser power= 0.28 mW/mm^2^).

**Figure S4.**
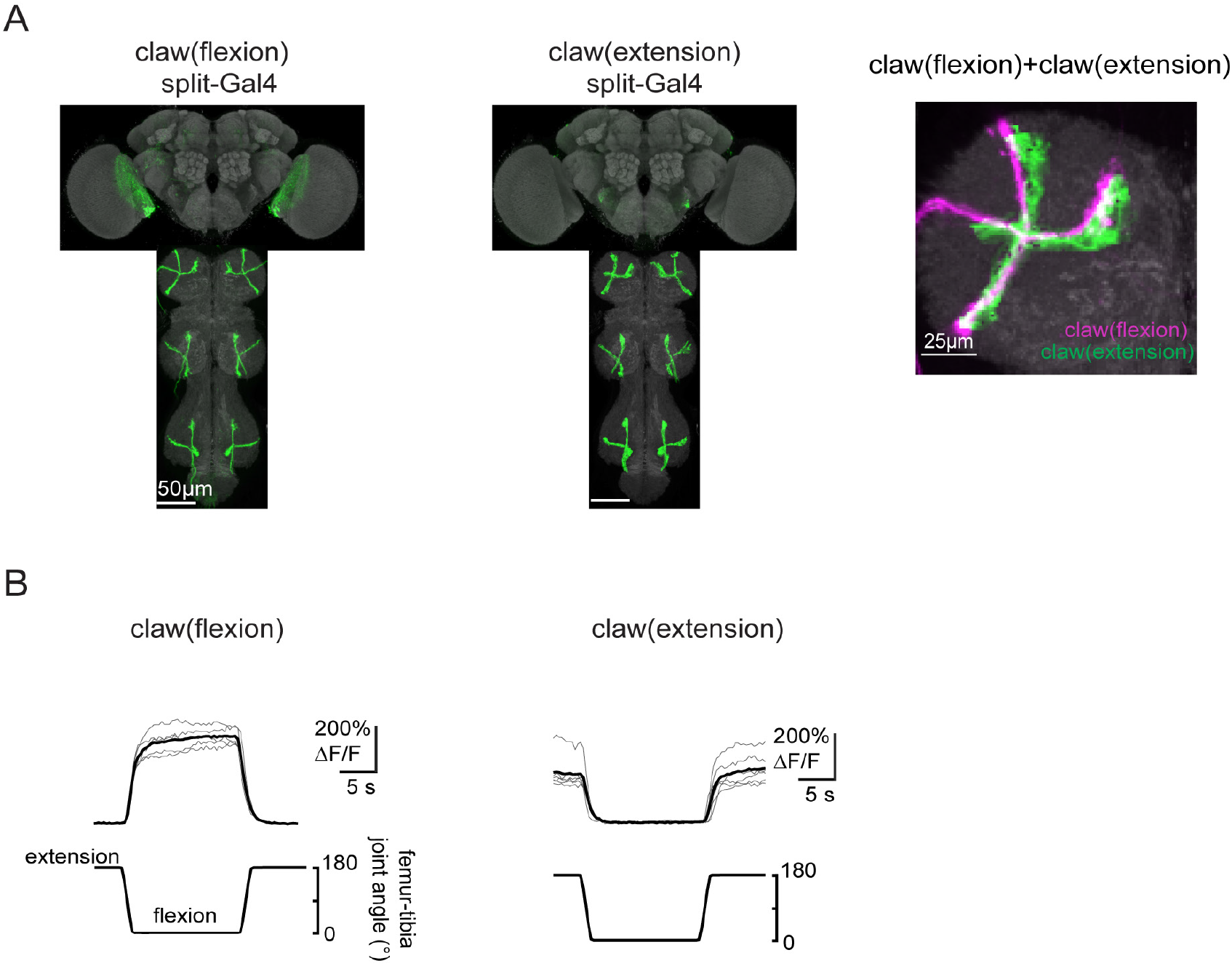
Distinct classes of claw neurons respond to tibia flexion and extension. **(A)** Genetic driver lines labeling claw neurons that encode tibia flexion and extension. GFP (green) expression in VNC and brain driven by split-Gal4 lines targeting subtypes of the claw neurons. Grey: neuropils and brains were stained with nc82. Right: co-localization of claw-flexion and claw-extension neurons. VNC images were aligned to a common template *in silico*. **(B)** Calcium responses of claw-flexion and claw-extension neurons during passive movement of the tibia (n=6 flies of each genotype).

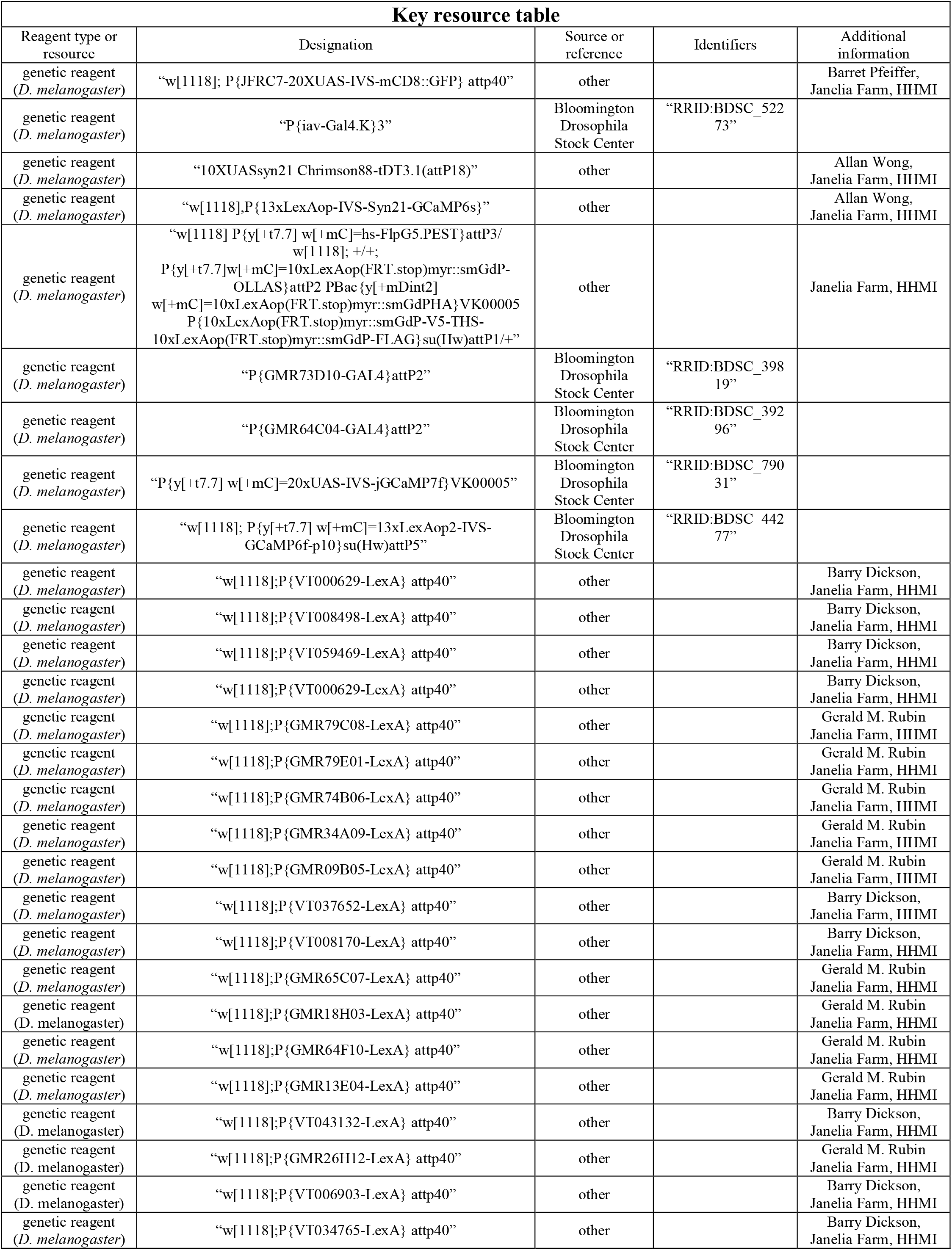
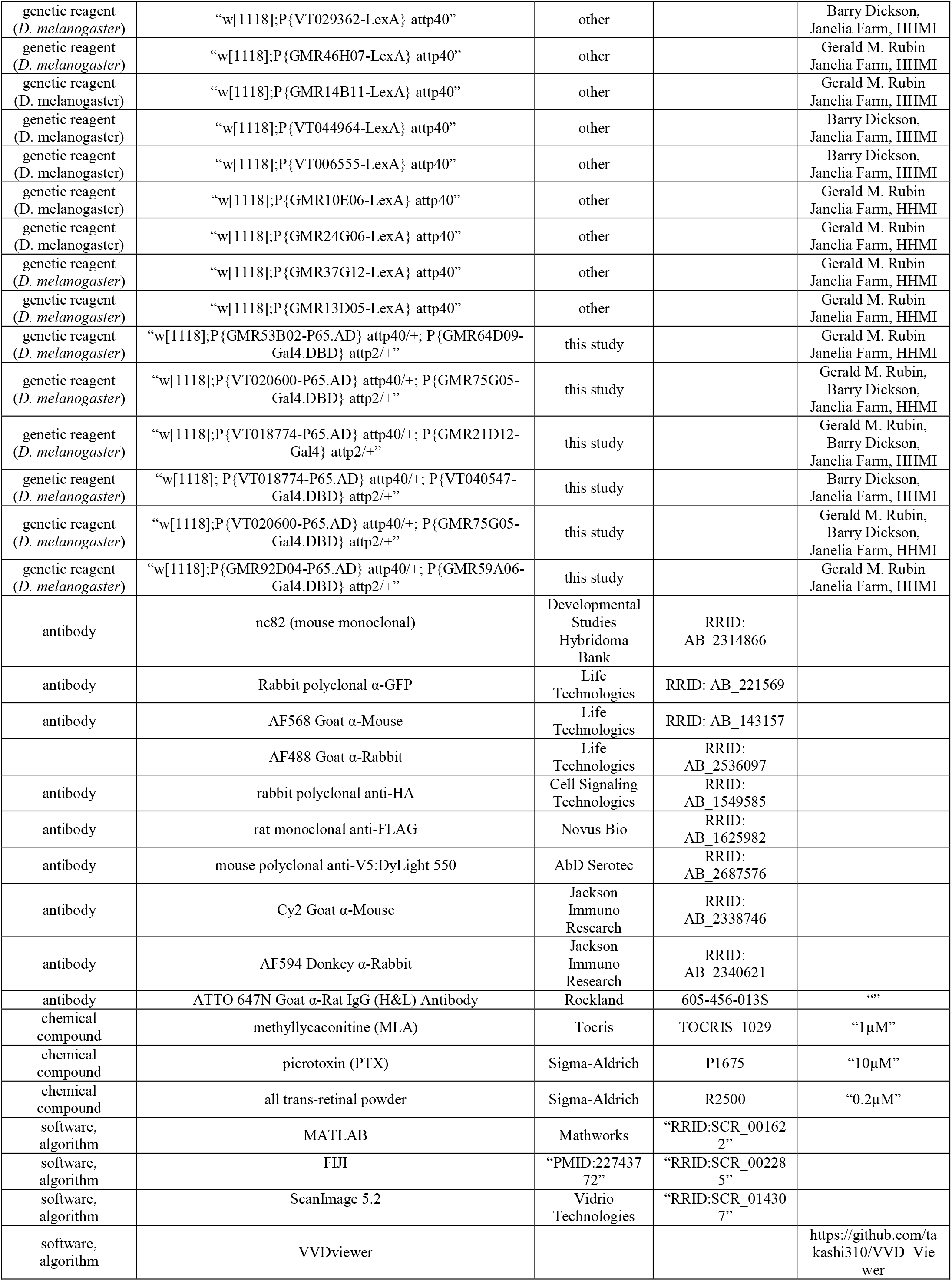

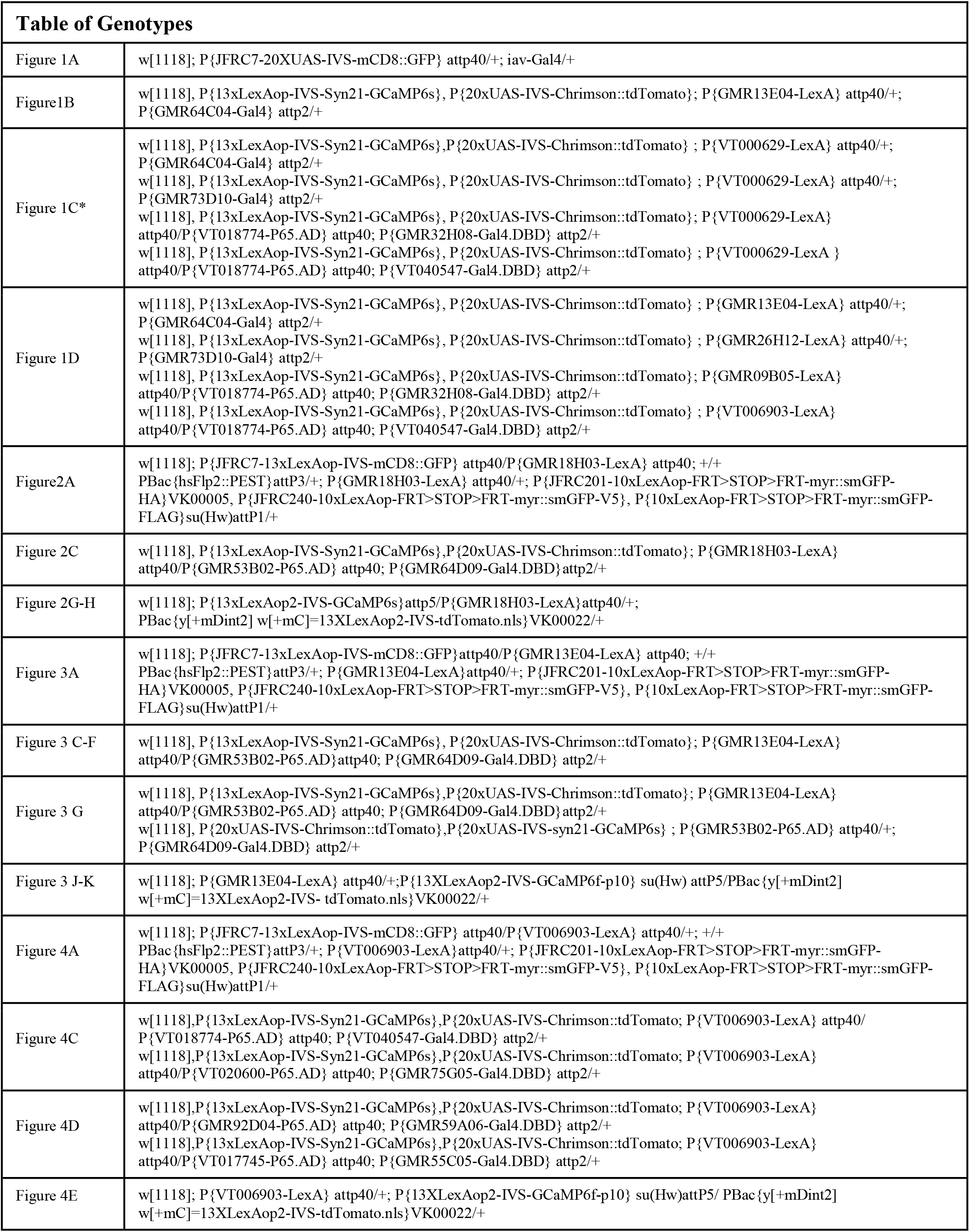
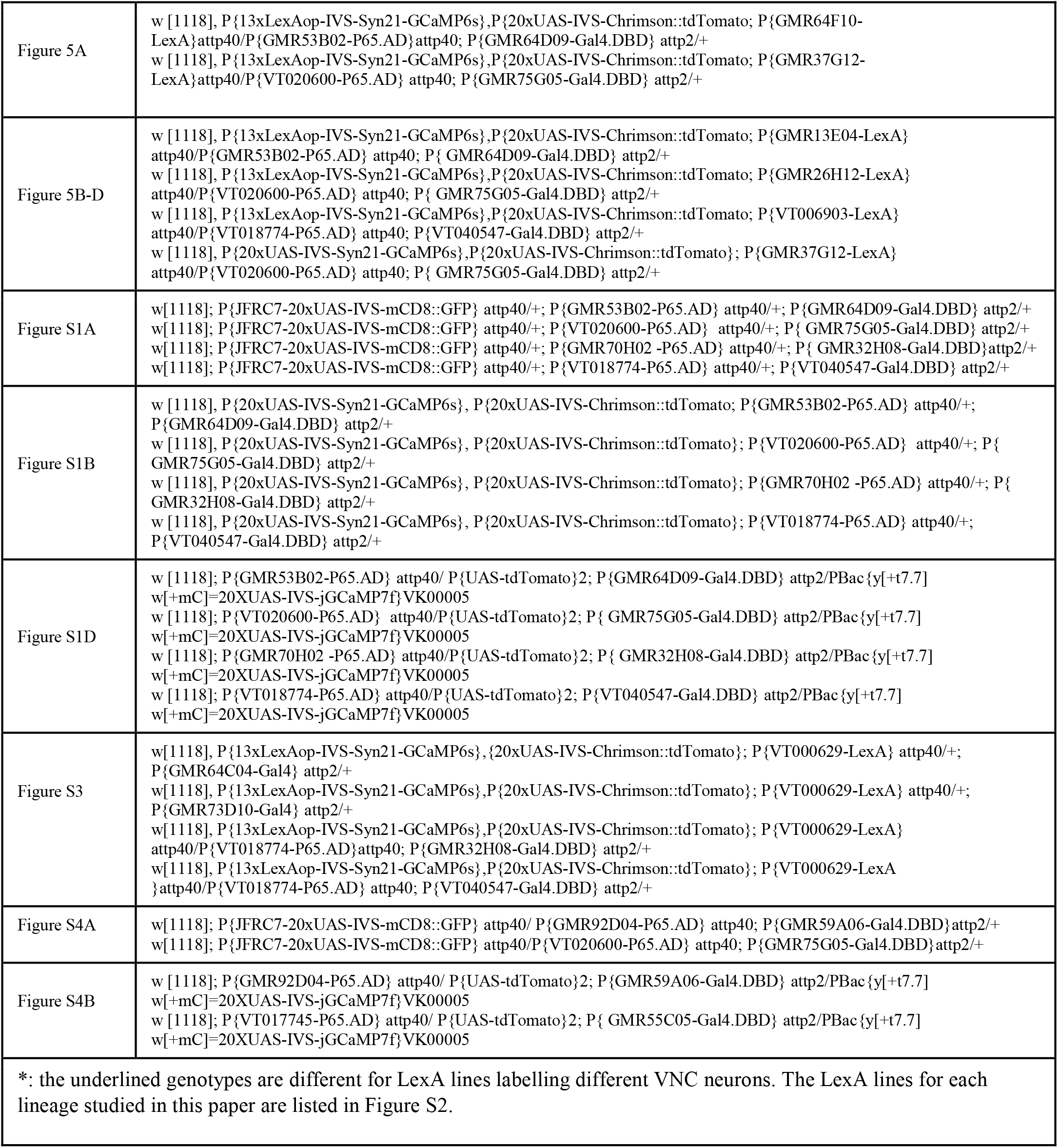

## References

Agrawal, S., Dickinson, E.S., Sustar, A., Gurung, P., Shepherd, D., Truman, J.W., and Tuthill, J.C. (2020). Central processing of leg proprioception in Drosophila. ELife 9, e60299.

Akay, T., Tourtellotte, W.G., Arber, S., and Jessell, T.M. (2014). Degradation of mouse locomotor pattern in the absence of proprioceptive sensory feedback. Proc Natl Acad Sci USA 111, 16877–16882.

Allen, A.M., Neville, M.C., Birtles, S., Croset, V., Treiber, C.D., Waddell, S., Goodwin, S.F., and Mann, R.S. (2020). A single-cell transcriptomic atlas of the adult Drosophila ventral nerve cord. ELife 9, 1–32.

Azevedo, A.W., Dickinson, E.S., Gurung, P., Venkatasubramanian, L., Mann, R.S., and Tuthill, J.C. (2020). A size principle for recruitment of Drosophila leg motor neurons. ELife 9, e56754.

Bässler, U. (1993). The femur-tibia control system of stick insects — a model system for the study of the neural basis of joint control. Brain Research Reviews 18, 207–226.

Bässler, U., and Büschges, A. (1998). Pattern generation for stick insect walking movements—multisensory control of a locomotor program. Brain Research Reviews 27, 65–88.

Bässler, U., Wolf, H., and Stein, W. (2007). Functional recovery following manipulation of muscles and sense organs in the stick insect leg. J Comp Physiol A 193, 1151–1168.

Bates, A.S., Janssens, J., Jefferis, G.S., and Aerts, S. (2019). Neuronal cell types in the fly: single-cell anatomy meets single-cell genomics. Current Opinion in Neurobiology 56, 125–134.

Bidaye, S.S., Bockemühl, T., and Büschges, A. (2018). Six-legged walking in insects: How CPGs, peripheral feedback, and descending signals generate coordinated and adaptive motor rhythms. Journal of Neurophysiology 119, 459–475.

Burns, M.D. (1974). Structure and physiology of the locust femoral chordotonal organ. Journal of Insect Physiology 20, 1319–1339.

Burrows, M. (1996). The Neurobiology of an Insect Brain-Motor Neurons. (Oxford: Oxford University press)

Büschges, A. (1989). Processing of Sensory Input from the Femoral Chordotonal Organ by Spiking Interneurones of Stick Insects. Journal of Experimental Biology 144, 81–111.

Cafaro, J., and Rieke, F. (2013). Regulation of spatial selectivity by crossover inhibition. J Neurosci 33, 6310– 6320.

Chen, T.-W., Wardill, T.J., Sun, Y., Pulver, S.R., Renninger, S.L., Baohan, A., Schreiter, E.R., Kerr, R.A., Orger, M.B., Jayaraman, V., et al. (2013). Ultrasensitive fluorescent proteins for imaging neuronal activity. Nature 499, 295–300.

Court, R., Namiki, S., Armstrong, J.D., Börner, J., Card, G., Costa, M., Dickinson, M., Duch, C., Korff, W., Mann, R., et al. (2020). A Systematic Nomenclature for the Drosophila Ventral Nerve Cord. Neuron 107, 1071-1079.e2.

Davis, F.P., Nern, A., Picard, S., Reiser, M.B., Rubin, G.M., Eddy, S.R., and Henry, G.L. (2020). A genetic, genomic, and computational resource for exploring neural circuit function (eLife Sciences Publications Limited).

Dionne, H., Hibbard, K.L., Cavallaro, A., Kao, J.-C., and Rubin, G.M. (2018). Genetic Reagents for Making Split-GAL4 Lines in Drosophila. Genetics 209, 31–35.

Dubs, A., Laughlin, S.B., and Srinivasan, M.V. (1981). Single photon signals in fly photoreceptors and first order interneurones at behavioral threshold. J Physiol 317, 317–334.

Fabre, C.C.G., Hedwig, B., Conduit, G., Lawrence, P.A., Goodwin, S.F., and Casal, J. (2012). Substrate-borne vibratory communication during courtship in Drosophila melanogaster. Curr Biol 22, 2180–2185.

Franconville, R., Beron, C., and Jayaraman, V. (2018). Building a functional connectome of the drosophila central complex. ELife 7, 1–24.

French, A.S., and Wong, R.K.S. (1976). The responses of trochanteral hair plate sensilla in the cockroach to periodic and random displacements. Biological Cybernetics 22, 33–38.

Grimes, W.N., Schwartz, G.W., and Rieke, F. (2014). The synaptic and circuit mechanisms underlying a change in spatial encoding in the retina. Neuron 82, 460–473.

Harris, R.M., Pfeiffer, B.D., Rubin, G.M., and Truman, J.W. (2015). Neuron hemilineages provide the functional ground plan for the Drosophila ventral nervous system. ELife 4, 1–34.

Hill, P.S.M., and Wessel, A. (2016). Biotremology. Current Biology 26, R187–R191.

Jenett, A., Rubin, G.M., Ngo, T.-T.B., Shepherd, D., Murphy, C., Dionne, H., Pfeiffer, B.D., Cavallaro, A., Hall, D., Jeter, J., et al. (2012). A GAL4-Driver Line Resource for Drosophila Neurobiology. Cell Reports 2, 991– 1001.

Klapoetke, N.C., Murata, Y., Kim, S.S., Pulver, S.R., Birdsey-Benson, A., Cho, Y.K., Morimoto, T.K., Chuong, A.S., Carpenter, E.J., Tian, Z., et al. (2014). Independent optical excitation of distinct neural populations. Nat Methods 11, 338–346.

Kuan, A.T., Phelps, J.S., Thomas, L.A., Nguyen, T.M., Han, J., Chen, C.-L., Azevedo, A.W., Tuthill, J.C., Funke, J., Cloetens, P., et al. (2020). Dense neuronal reconstruction through X-ray holographic nano-tomography. Nat Neurosci 23, 1637–1643.

Lacin, H., and Truman, J.W. (2016). Lineage mapping identifies molecular and architectural similarities between the larval and adult Drosophila central nervous system. ELife 5, 1–28.

Lacin, H., Chen, H.M., Long, X., Singer, R.H., Lee, T., and Truman, J.W. (2019). Neurotransmitter identity is acquired in a lineage-restricted manner in the Drosophila CNS. ELife 8, 1–26.

Liu, W.W., and Wilson, R.I. (2013). Glutamate is an inhibitory neurotransmitter in the Drosophila olfactory system. 10.

Liu, W.W., Mazor, O., and Wilson, R.I. (2015). Thermosensory processing in the Drosophila brain. Nature 519, 353–357.

Mamiya, A., Gurung, P., and Tuthill, J.C. (2018). Neural Coding of Leg Proprioception in Drosophila. Neuron 100, 636-650.e6.

Matheson, T., and Field, L.H. (1995). An elaborate tension receptor system highlights sensory complexity in the hind leg of the locust. Journal of Experimental Biology 198, 1673–1689.

Mendes, C.S., Bartos, I., Akay, T., Márka, S., and Mann, R.S. (2013). Quantification of gait parameters in freely walking wild type and sensory deprived Drosophila melanogaster. ELife 2, e00231.

Morimoto, M.M., Nern, A., Zhao, A., Rogers, E.M., Wong, A.M., Isaacson, M.D., Bock, D.D., Rubin, G.M., and Reiser, M.B. (2020). Spatial readout of visual looming in the central brain of Drosophila. ELife 9, e57685.

Namiki, S., Dickinson, M.H., Wong, A.M., Korff, W., and Card, G.M. (2018). The functional organization of descending sensory-motor pathways in drosophila. ELife 7, 1–50.

Nern, A., Pfeiffer, B.D., and Rubin, G.M. (2015). Optimized tools for multicolor stochastic labeling reveal diverse stereotyped cell arrangements in the fly visual system. Proc Natl Acad Sci U S A 112, E2967–2976.

Otsuna, H., Ito, M., and Kawase, T. (2018). Color depth MIP mask search: a new tool to expedite Split-GAL4 creation. BioRxiv 318006.

Patella, P., and Wilson, R.I. (2018). Functional Maps of Mechanosensory Features in the Drosophila Brain. Current Biology 28, 1189-1203.e5.

Phelps, J.S., Hildebrand, D.G.C., Graham, B.J., Kuan, A.T., Thomas, L.A., Nguyen, T.M., Buhmann, J., Azevedo, A.W., Sustar, A., Agrawal, S., et al. (2021). Reconstruction of motor control circuits in adult Drosophila using automated transmission electron microscopy. Cell.

Prsa, M., Morandell, K., Cuenu, G., and Huber, D. (2019). Feature-selective encoding of substrate vibrations in the forelimb somatosensory cortex. Nature 567, 384–388.

Scheffer, L.K., Xu, C.S., Januszewski, M., Lu, Z., Takemura, S.-Y., Hayworth, K.J., Huang, G.B., Shinomiya, K., Maitlin-Shepard, J., Berg, S., et al. (2020). A connectome and analysis of the adult Drosophila central brain. Elife 9.

Schindelin, J., Arganda-Carreras, I., Frise, E., Kaynig, V., Longair, M., Pietzsch, T., Preibisch, S., Rueden, C., Saalfeld, S., Schmid, B., et al. (2012). Fiji: an open-source platform for biological-image analysis. Nature Methods 9, 676–682.

Schneider-Mizell, C.M., Gerhard, S., Longair, M., Kazimiers, T., Li, F., Zwart, M.F., Champion, A., Midgley, F.M., Fetter, R.D., Saalfeld, S., et al. (2016). Quantitative neuroanatomy for connectomics in Drosophila. ELife 5, e12059.

Shepherd, D., Sahota, V., Court, R., Williams, D.W., and Truman, J.W. (2019). Developmental organization of central neurons in the adult Drosophila ventral nervous system. J Comp Neurol 527, 2573–2598.

Siegler, M.V.S., and Burrows, M. (1983). Spiking Local Interneurons as Primary Integrators of Mechanosensory Information in the Locust. 15.

Stein, W., and Sauer, A.E. (1999). Physiology of vibration-sensitive afferents in the femoral chordotonal organ of the stick insect. Journal of Comparative Physiology A: Sensory, Neural, and Behavioral Physiology 184, 253– 263.

Takeoka, A., Vollenweider, I., Courtine, G., and Arber, S. (2014). Muscle Spindle Feedback Directs Locomotor Recovery and Circuit Reorganization after Spinal Cord Injury. Cell 159, 1626–1639.

Tirian, L., and Dickson, B.J. (2017). The VT GAL4, LexA, and split-GAL4 driver line collections for targeted expression in the Drosophila nervous system. BioRxiv 198648.

Truman, J.W., Moats, W., Altman, J., Marin, E.C., and Williams, D.W. (2010). Role of Notch signaling in establishing the hemilineages of secondary neurons in Drosophila melanogaster. Development 137, 53–61.

Tuthill, J.C., and Azim, E. (2018). Proprioception. Current Biology 28, R194–R203.

Tuthill, J.C., and Wilson, R.I. (2016a). Parallel Transformation of Tactile Signals in Central Circuits of Drosophila. Cell 164, 1046–1059.

Tuthill, J.C., and Wilson, R.I. (2016b). Mechanosensation and Adaptive Motor Control in Insects. Current Biology 26, R1022–R1038.

Venkatasubramanian, L., and Mann, R.S. (2019). The development and assembly of the Drosophila adult ventral nerve cord. Current Opinion in Neurobiology 56, 135–143.

Windhorst, U. (2007). Muscle proprioceptive feedback and spinal networks. Brain Research Bulletin 73, 155– 202.

Zheng, Z., Lauritzen, J.S., Perlman, E., Robinson, C.G., Nichols, M., Milkie, D., Torrens, O., Price, J., Fisher, C.B., Sharifi, N., et al. (2018). A Complete Electron Microscopy Volume of the Brain of Adult Drosophila melanogaster. Cell 174, 730-743.e22.

Zill, S., Schmitz, J., and Büschges, A. (2004). Load sensing and control of posture and locomotion. Arthropod Structure & Development 33, 273–286.

